# Learning from ingroup experiences changes intergroup impressions

**DOI:** 10.1101/2021.11.02.466926

**Authors:** Yuqing Zhou, Björn Lindström, Alexander Soutschek, Pyungwon Kang, Philippe N. Tobler, Grit Hein

**Author notes:** contributed equally to this work (Yuqing Zhou, Björn Lindström). shared senior authorship (Philippe N. Tobler, Grit Hein). Address correspondence to: Prof. Dr. Grit Hein, Translational Social Neuroscience Lab, University Hospital of Würzburg, Margarete-Höppel-Platz 1, 97080 Würzburg, Germany, Dr. Yuqing Zhou, Translational Social Neuroscience Lab, University Hospital of Würzburg, Margarete-Höppel-Platz 1, 97080 Würzburg, Germany.

## Abstract

Living in multicultural societies, humans form impressions towards individuals of their own social groups (ingroup members) and of different social groups (outgroup members). Some psychological theories predict that intergroup impressions are mainly shaped by experiences with outgroup individuals (“outgroup focused theories”), while other theories predict that ingroup experiences play a dominant role (“ingroup focused theories”). Here we test predictions from these two psychological theories by estimating how intergroup impressions are dynamically shaped when people learn from both ingroup and outgroup experiences. Participants expected to receive painful shocks but were saved from pain by different ingroup or outgroup members in 75% of all trials. We measured neural responses using functional magnetic resonance imaging (fMRI), and participants rated their social closeness as well as impressions towards the ingroup and the outgroup. Behavioral results showed an initial ingroup bias in impression ratings which was significantly reduced over the course of learning. Computational learning models revealed that these changes in intergroup impressions were predicted by the weight given to ingroup prediction errors. The weight of the ingroup prediction error and its effect on intergroup impression change was stronger the more individuals identified with their ingroup. On the neural level, the left inferior parietal lobule (IPL) encoded more negative prediction errors for the ingroup compared to the outgroup. Moreover, stronger weight for ingroup prediction errors was related to increased neural coupling between the left IPL and the anterior insula (AI). This coupling further predicted learning-related changes in intergroup impressions. Together, our work provides computational and neural evidence for “ingroup focused theories”, highlighting the importance of ingroup experiences in shaping social impressions in intergroup settings.

## Introduction

In multicultural societies, people encounter individuals from their own social group (ingroup members) and from different social groups (outgroup members), and form impressions towards these ingroup and outgroup individuals. As impressions predict behaviors (Tajfel et al., 1971; Greenwald et al., 2009; Amodio & Cikara, 2021), it is important to understand the mechanisms that shape impressions in intergroup contexts.

One basic mechanism that shapes impressions is learning (Hein et al., 2016a; Mende-Siedlecki, 2018; Siegel et al., 2018; Hackel et al., 2015; Tobler et al., 2006; Burke et al., 2010; Kang et al., 2020; Lindström et al., 2014; Will et al., 2017). According to learning theory, positive experiences with a person establish positive associations towards this individual, leading people to like them, to interact with them and to reciprocate (Hackel et al., 2019; Hackel & Zaki, 2018; Pettigrew & Tropp, 2006; Lindström & Tobler, 2018). Negative experiences have the opposite effect: they establish negative associations, which give rise to negative impressions (Barlow et al., 2012). Moreover, unpredicted positive or negative experiences elicit prediction errors, i.e. discrepancies between predictions and experienced outcomes (Rescorla & Wagner, 1972), and the updating of impressions regarding others corresponds to gradually reducing prediction errors by adjusting predictions about them (Mende-Siedlecki, 2018; Hackel & Amodio, 2018). Importantly, in complex social environments, individuals learn from intermittent experiences with both ingroup and outgroup members. However, it remains unclear how such ingroup and outgroup experiences dynamically shape neural learning processes, and learning-related changes in impressions towards in- and outgroups.

One class of influential social science theories proposes that intergroup impressions are mainly shaped by experiences with outgroup individuals (“outgroup focused theories”). Previous evidence showed that positive outgroup contact decreased ingroup-outgroup boundaries (Dovidio et al., 2003; Bettencourt et al., 1992; Marcus-Newhall et al., 1993), reduced outgroup prejudice (Pettigrew & Tropp, 2006; Schlueter & Scheepers, 2010; Chu & Griffey, 1985), and increased empathy for outgroup members (Paluck, 2009; Malhotra & Liyanage, 2005; Hein et al., 2016a). Furthermore, expectations concerning outgroup members are typically more negative than expectations concerning ingroup members (Dovidio & Gaertner, 1993). From a learning theory perspective, it is plausible to assume that positive contact with an outgroup individual is an unexpected positive outcome and may lead to stronger positive prediction errors compared with positive ingroup experiences. Consistent with this assumption, previous studies showed an increase in outgroup positivity after positive outgroup experiences, while the same positive experiences with the ingroup had no significant effect (Hein et al., 2016a; 2018). On the neural level, learning from outgroup experiences was associated with activation in the anterior insular cortex (Hein et al., 2016a).

Alternatively, another class of social science theories proposes that changes of intergroup impressions are mainly driven by ingroup experiences (“ingroup focused theories”). It is based on the assumption that maintaining a positive self-image requires a positive social identity, which in turn requires a more positive evaluation of ingroup compared to outgroup individuals (Tajfel et al., 1979). A frequent consequence of such differential evaluations are ingroup biases, i.e. more positive impressions towards ingroup compared to outgroup individuals (Tajfel & Turner, 1979; Maass et al., 1989; Foddy et al., 2009), and a reluctance to process negative ingroup information (Hughes et al., 2017; Howard & Rothbart, 1980; Maass et al., 1989). These ingroup biases (fueled by the need to maintain a positive social identity) are challenged if ingroup individuals deviate from ingroup norms or expectations, because unexpected negative experiences with ingroup individuals threaten the positive demarcation from the outgroup (Hutchison et al., 2008; Marques et al., 1988; Mendoza et al., 2014). The effect is enhanced in individuals that strongly identify with their ingroup (Biernat et al., 1999; Branscombe et al., 1993; Mendoza et al., 2014). In sum, according to “outgroup focused theories”, changes in intergroup impressions should be mainly driven by learning from unexpected outgroup experiences, while “ingroup focused theories” predict that changes in intergroup impressions are mainly driven by unexpected ingroup experiences.

Previous studies have investigated how learning from unexpected experiences (prediction errors) shapes impression formation towards individuals of the same group, i.e., in the absence of a social group manipulation (Hackel et al., 2015; Ligneul et al., 2016; Park et al., 2021). At the neural level, these studies associated learning-related updates of impressions with changes in a neural network, centered on medial prefrontal cortex (mPFC), inferior parietal lobule (IPL), ventrolateral prefrontal cortex (vlPFC), posterior cingulate cortex (PCC), striatum and the temporoparietal junction (TPJ). These results have provided first insights into the neural circuitries that link reinforcement learning and impression formation. However, it remained unclear whether and how learning from intermittent experiences with ingroup and outgroup individuals, the type of experience that is most natural in complex social systems and modern multicultural societies (Berry & Sam, 2014), changes intergroup impressions and underlying neural circuitries.

To address this question, we designed an experiment in which identical experiences with the in- and outgroup could dynamically affect the impressions towards both groups. In our experiment, participants initially met members of their own group (six fellow Swiss participants; ingroup) and of a different group (six Middle Eastern confederates; outgroup) (see Methods for the pre-scanning procedure). The two groups were seated in adjacent rooms (i.e., an “ingroup room” and an “outgroup room”), denoted as “Room A” and “Room B”, counterbalanced across participants.

The participant inside the scanner underwent a modified version of a Pavlovian learning experiment involving pain stimulation. In each trial an individual from the ingroup or the outgroup (i.e., from room A or B) ostensibly could give up money to save the participant from pain. In reality, participants were relieved from pain in 75% of all trials based on a predefined algorithm. Importantly, this ratio was the same in the ingroup and the outgroup condition. Thus, objective experiences with the two groups were equalized, and primarily positive.

In the beginning of each trial, participants used a manikin representing themselves to move towards or away from the ingroup (e.g., Room “A”) or the outgroup (e.g., Room “B”; **Figure 1**), providing a measure of ingroup and outgroup closeness that could change through learning. Next, participants were presented a cue (letter), indicating the room in which the interaction partner would sit. Note that the letter only indicated a room and not a specific ingroup or outgroup person. This is important, because it allowed us to assess how experiences with ingroup and outgroup individuals shape closeness and impressions toward the entire ingroup and outgroup, instead of specific ingroup and outgroup individuals. Following the room cue, participants rated the degree to which they expected to receive pain. Then, the ostensible decision of the other person was revealed, and implemented (**Figure 1**). Before and after the experiment, participants provided ingroup and outgroup impressions on a questionnaire (Hein et al., 2016a). Moreover, prior to the learning experiment, we assessed the degree to which participants identified with the ingroup (ingroup identification questionnaire, Leach et al., 2008), and disliked the outgroup (modified version of the Modern Racism Scale, McConahay, 1986). This setup allowed us to model how participants learn from the same in- and outgroup experiences and how these learning processes shape intergroup impressions on the behavioral and neural level.

**Figure 1.**
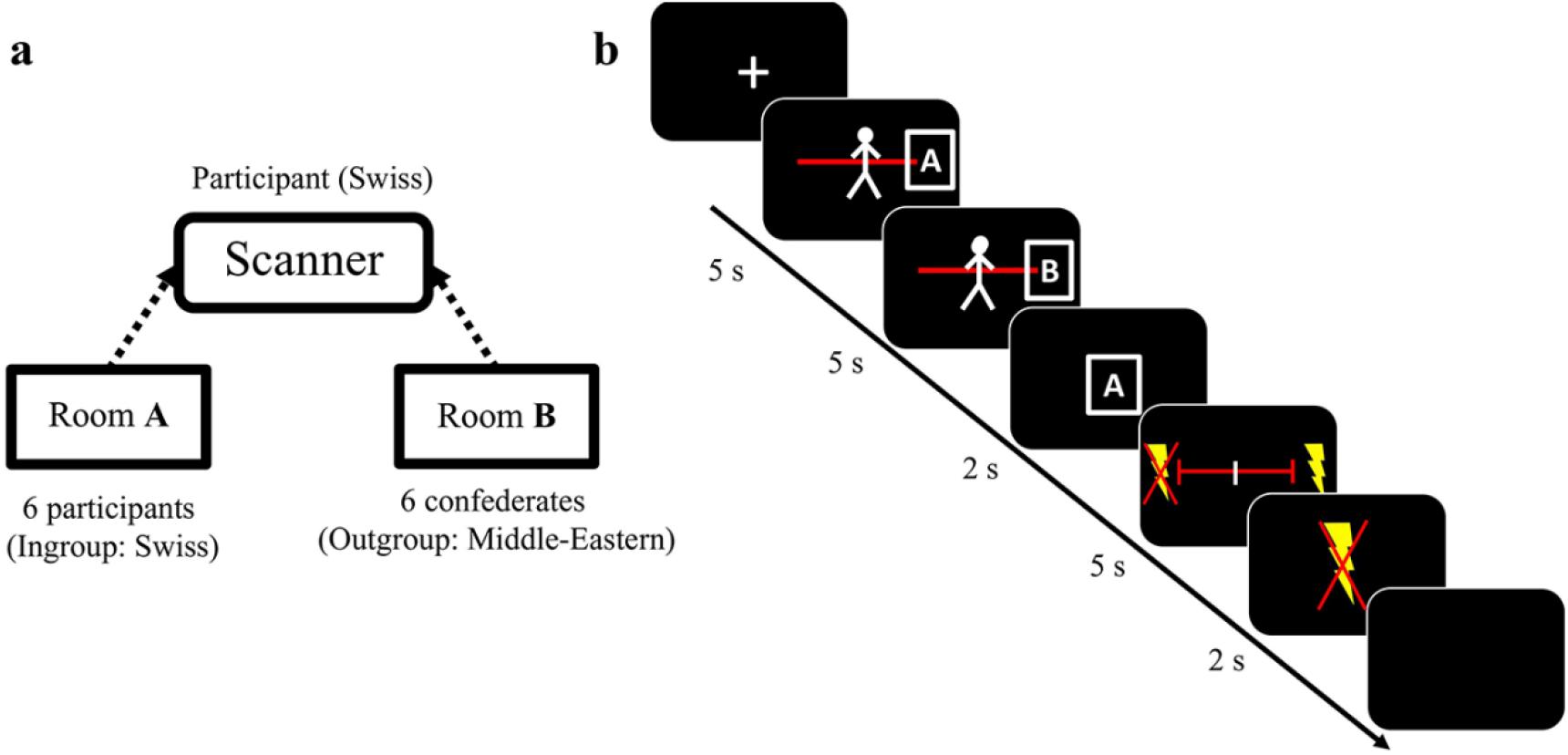
Experimental task. (a) Experimental procedure. The Swiss participant in the scanner interacted with both ingroup members (six Swiss participants, in this example seated in Room A) and outgroup members (six Middle Eastern confederates, in this example seated in Room B) during the experiment. The assignment of room type (A or B) to the respective social groups (ingroup or outgroup) was counterbalanced across participants. (b) Example trial sequence. At the beginning of each trial, the participants rated their closeness towards the in- and outgroup on separate rating scales that were presented in randomized order. To do so, they moved a manikin representing themselves towards or away from the room in which ingroup or outgroup members were seated. The next screen revealed the room in which the person that could influence the participant’s pain was located (e.g., room “A”, which in this example represented the ingroup). Next, participants rated their expectancy of receiving pain, followed by the symbol that indicated the ostensible decision of the other person (a crossed-out lightning bolt representing the decision to give up money to save the participant from pain and a lightning bolt representing the decision to keep the money, resulting in pain for the participant). The experienced outcome (pain or no pain) was delivered during the final blank screen.

Based on previous evidence (Dovidio & Gaertner, 1993), we hypothesized that initially participants would feel closer and show more positive impressions towards the ingroup compared to the outgroup. Based on the implications of learning theory (Rescorla & Wagner, 1972; Hein et al., 2016a), we anticipated that equal experiences with the ingroup and outgroup would result in a decreased group difference in impressions. According to outgroup focused theories, the effect should be driven by outgroup prediction errors, i.e., prediction errors that are elicited by the unexpected experiences with the outgroup members (e.g., being-saved from pain in most trials; Hein et al., 2016a, 2018). In this case, learning from outgroup prediction errors, captured by trial-by-trial changes in closeness ratings, should predict the ingroup vs. outgroup impression difference after learning. In contrast, according to ingroup focused theories, changes in intergroup impressions should be linked to ingroup prediction errors, i.e., prediction errors that are elicited by unexpected experiences with ingroup members (i.e., not being saved from pain in some trials), with stronger effects in individuals who identify more strongly with the ingroup. In this case, the ingroup vs. outgroup impression difference after learning should be predicted by ingroup prediction errors, captured by respective changes in trial-by-trial closeness ratings.

On the neural level, the respective learning processes should be related to activations in the brain regions that have been associated with the updating of impressions (Mende-Siedlecki et al., 2013, Mende-Siedlecki, 2018, Hackel et al., 2015, Ligneul et al., 2016; Park et al., 2021; Hughes et al., 2017). These regions include the mPFC, IPL, vlPFC, PCC, striatum, and TPJ. The resulting change in the ingroup vs. outgroup difference in impression may be tracked by the anterior insula, given that this region has been linked to increased outgroup positivity (Hein et al., 2016a; Sheng et al., 2014; Wang et al., 2015).

## Methods

### Participants

30 healthy males (mean age ± SD = 22.7 ± 2.8 years) participated in this study as paid volunteers. We chose an all-male instead of a gender-mixed group of participants so that we could also use all-male confederates and avoid the complications of gender-mixed pairing of participants and confederates. All participants were born and raised in Switzerland and had normal or corrected-to-normal vision. The ingroup consisted of six additional male Swiss participants who eventually completed an independent study. The outgroup comprised six male confederates whose families originated from the Middle East. It was made clear that the individuals present in any given session did not know each other. One participant completed the fMRI study, but due to technical problems, only the behavioral results but not the imaging data was properly recorded. We report the behavioral and computational modelling results of 30 participants (mean age ± SD = 22.7 ± 2.8 years) in the main manuscript and the comparable behavioral results of 29 participants (mean age ± = 22.8 ± 2.8 years) in the supplementary material (**Table S4 – Table S9**). The behavioral and computational modelling findings were robust to the exclusion of the one participant (see **Table S4 – Table S9**). The imaging results are based on the available 29 imaging data sets. A sensitivity analysis using G*Power 3.1 indicates that given α (5%) and considering 4 predictors in the regression model, the current sample size (N = 30) has 80% power to detect a moderate effect size of f^2^ = 0.28 and critical t = 2.06. The study was approved by the Research Ethics Committee of the Canton (state) of Zurich. All participants provided written informed consent after the experimental procedure had been fully explained. Participants were reminded of their right to withdraw at any time during the study.

### Prescanning procedure

At the beginning of the testing session, an experimenter welcomed both the Swiss participants and the Middle Eastern confederates and informed them that the aim of the experiment was to measure cultural differences in decision making. After the introduction, all test participants were assigned a number ranging from 1 to 13 which determined their place in the testing lab. The testing lab consisted of two rooms that could be separated by a sliding door. Care was taken to ensure that all Swiss participants and all Middle East confederates were placed in different partitions. The separated rooms were named “Room A” and “Room B”, and the relationships between the room name and group membership were counterbalanced across participants.

After the room assignment, a well-established priming procedure (Dijksterhuis & van Knippenberg, 1998) served to enhance the social group manipulation. Specifically, participants were asked to write down five attributes that are typical for Middle Eastern males and for Swiss males. This procedure is commonly used to activate group-related stereotypes (Hein et al., 2016a), in our case to activate the stereotype of Middle Eastern males in our Swiss participants.

The six Middle Eastern confederates and six of the Swiss participants remained in Room A and Room B. The seventh Swiss participant that had signed up for the fMRI study was taken to the control room of the fMRI scanner. Here, a pain electrode was attached to the back of the left hand of the participant, and the individual pain threshold was determined based on a standard procedure (Hein et al., 2016a and 2016b). In particular, the participant was asked to rate the intensity of the electrical stimulation he received from 1 (not painful) to 10 (extremely painful). We used the intensity corresponding to a subjective value of 8 as the intensity inside the scanner. Electrical stimulation (bipolar, monophasic; maximum duration: 1,000 ms; input range, 5 V; output range, 50 mA) was delivered with a single-current stimulator (DS5; Digitimer, Ltd). Before entering the fMRI scanner, the participant was informed that he might receive painful stimulation, and in each trial, one of the individuals in Room A or Room B could prevent the upcoming painful stimulation for the participant by giving up money he would otherwise earn.

To avoid possible reputation effects that might influence the behavior of our participants, all ratings were kept anonymous, and we ensured that the participants did not meet the confederates after the experiment. Thus, all ratings and decisions were personal and could not be observed by the other participants. Everyone was also clearly informed that they would not meet after the experiment because the scanned participant needed to stay longer for an anatomical scan.

### Scanning Procedure

During the fMRI scans, each trial began with a fixation cross presented for 1-7 seconds. Next, the first closeness rating scale was shown for 5 s, followed by the second closeness rating scale, also presented for 5 s. For both ratings, participants moved a manikin that represented themselves on a 10-step scale towards or away from Room A or Room B to indicate how close they felt towards the individuals in the respective room. To avoid movement preparation and to keep the participants engaged, the manikin was placed at a random location on the scale. The order of closeness ratings for Rooms A and B was randomized across trials.

After the closeness ratings, a letter was shown for 2 s, indicating the room in which the interaction partner (i.e., the person that could ostensibly save the participant from pain) was located. Next, the 10-step rating scale was presented for 5 s and participants rated their expectancy of receiving pain in the current trial from not likely at all (indicated by a crossed-out lightning bolt) to very likely (indicated by a lightning bolt). To maintain the attention of the participant and reduce motion preparation in each trial, the starting point of the rating scales occurred at a random position. Thereafter, the decision of the other individual was shown for 2 s. A crossed-out lightning bolt symbol indicated that one of the individuals from the respective room had given up 5 CHF to save the participant from pain, whereas a lightning bolt symbol indicated upcoming painful stimulation. Based on the decision, participants then either did or did not receive painful stimulation during the final blank screen.

Participants were relieved from pain in 75% of all trials. Because this ratio was the same in the ingroup and the outgroup condition, the objective experiences with the two groups were equalized, and primarily positive. The learning experiment consisted of three blocks, with 24 trials each, resulting in 72 trials in total.

### Questionnaires

We used an impression scale to assess participants’ impressions towards ingroup (Swiss) and outgroup (Middle Eastern) individuals. Participants rated their impressions towards individuals in Room A and Room B separately before and after the learning experiment. The impression scale included questions regarding perceived group membership (e.g. “To what extent do you see yourself and these other persons as part of the same group”), similarity (e.g., “How much do you think you and these persons have in common?”), and likability (“How likable do you find these persons?”) (Hein et al., 2010, 2016a, 2018). Moreover, participants completed the ingroup identification questionnaire (Leach et al., 2008), which measures identification and satisfaction with the ingroup in general. Finally, participants indicated their tendency for outgroup discrimination using the Modern Racism Scale, modified to the Middle Eastern context (McConahay, 1986).

### Analyses of impression and closeness ratings

We performed linear mixed models (LMM, ‘lme4’, Bates et al., 2015) in R-3.5.1 (R Development Core Team, 2012) for the behavioral analyses on impression and closeness ratings. In particular, we conducted LMMs with group (ingroup/outgroup), time (before/after-learning) and group × time as predictors, and the impression/closeness rating scores as the dependent variable. We used by-participant random intercepts and slopes for all fixed effects. Likelihood ratio tests (LMT) were applied to assess the significance of the fixed effects. The resulting *χ^2^* values indicate how much more likely the data are under the assumption of a more complex model (i.e. a model including a particular parameter) than under the assumption of a simpler model (i.e. a model not including this particular parameter).

Individual ingroup and outgroup impression scores were calculated based on the average of the scores for each item of the respective ingroup or outgroup impression scales that participants completed before and after the learning experiment. The closeness ratings of three participants contained null values, which were replaced by the average closeness rating of all other participants at the corresponding trial.

### Computational modeling

#### Computational modeling of shock expectancy ratings

First, we modeled participants’ trial-by-trial shock expectancy ratings using a standard Rescorla-Wagner (Rescorla & Wagner, 1972) reinforcement learning (RL) algorithm. The RL model assumes that participants change their expectancy of being shocked when new information reveals that the experienced outcome is different from the expected outcome.

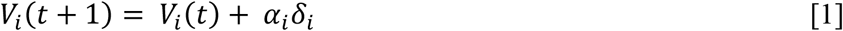

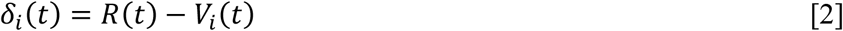

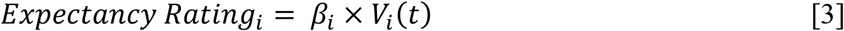

Therefore, on each trial *t*, the value of the (future) expectation *V_i_*(*t* + 1) for group *i* (ingroup or outgroup) is a function of current value *V_i_*(*t*) and the prediction error *δ_i_* (Equation 1), which corresponds to the difference between the experienced outcome *R*(*t*) at trial *t* (coded as 0 or 1 for shock or no shock) and the current value *V_i_* (Equation 2). Finally, the current value is multiplied by a response parameter *β* (*β* > 0), which linearly maps the current value to expectancy ratings (Equation 3). The learning rate α (0 ≤ *α* ≤ 1) controls the extent to which the current shock expectancy is updated by new information. Consequently, a low learning rate corresponds to a slow integration of prediction errors into current values.

We fitted participants’ shock expectancy to several models, including models that assumed the same learning rate and response parameter (Learning Model 1), different learning rates and same response parameters (Learning Model 2), different learning rates and response parameters (Learning Model 3) for the ingroup and the outgroup, and different learning rates for positive and negative prediction errors generated by the ingroup or outgroup (Learning Model 4, four learning rates). The winning model (Learning Model 3) included different learning rates and response parameters for experiences with group *i* (ingroup or outgroup)

#### Computational modeling of closeness ratings

Next, we fitted the trial-by-trial closeness ratings regarding the ingroup or outgroup as a linear function of previous prediction errors, i.e., based on the winning model (Learning Model 3) determined above. As the closeness ratings tended to be auto-correlated between trials, we used the trial-wise difference in closeness ratings to reduce this correlation (a standard method in time series analysis, Seth, 2010). Notably, these trial-by-trial changes in closeness to the in- and outgroup were only weakly correlated (average *r* across subjects = 0.14). In all models, we assumed that changes in closeness to group *i* (i.e., the ingroup or the outgroup) were a linear function of the time-discounted sum of previous prediction errors to outcomes from *i* (as originating from the RL model, Equations 1-3).

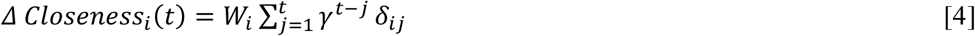

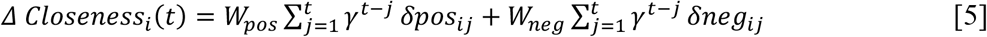

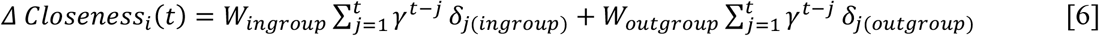

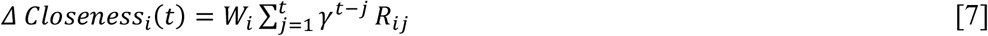

The winning model (Equation 4, Closeness Model 1) assumes closeness updates to group *i* to result from previous prediction errors to outcomes arising from *i*, and not to spill over between groups (e.g., receiving a shock from the outgroup did not predict a subsequent change in ingroup closeness). This model has two parameters. The first parameter *W* captures the magnitude (weight) of influence of prediction errors on changes in closeness. The *W*-parameter ranges from −10 to 10, because ten represents the maximum of the closeness ratings. Larger *W* corresponds to stronger influence of prediction errors on closeness updates. The second parameter is the discount parameter *γ* (0 ≤ *γ* ≤ 1), capturing an exponential decay of the influence of previous prediction errors over time, such that the more recent prediction errors have greater impact on the changes in closeness than the earlier prediction errors. If *γ* is close to one, all preceding prediction errors receive the same weight, and if it is close to zero, only the last prediction error leads to subsequent changes in closeness.

Closeness Model 2 (Equation 5) is similar to Closeness Model 1, except that we divided the prediction errors by experienced outcome and added them up separately. Accordingly, the model has three free parameters: weight for positive prediction errors from the corresponding group (*W_pos_*), weight for negative prediction errors from the corresponding group (*W_neg_*) and discounting parameter (*γ*). Closeness Model 3 (Equation 6) assumes that unexpected outcomes for the ingroup or the outgroup will lead to a change of closeness to both groups (e.g., receiving a shock from the outgroup could predict a subsequent change in ingroup closeness). This model has also three free parameters, weight for the ingroup prediction errors (*W_ingroup_*), weight for the outgroup prediction errors (*W_outgroup_*) and discounting parameter (*γ*). Finally, we fitted Closeness Model 4 (Equation 7), in which the change of closeness was determined solely by experienced (actual) outcome (*R*) to verify that the closeness update depends not only on the experience of shock, but on prediction errors. Equation 7 is similar to Equation 4 except that the sum of previous prediction errors has been replaced with the sum of previous outcome values.

#### Parameter estimation and model comparisons

To fit the parameters of the different computational models, we used a maximum likelihood approach, which determines the set of parameters that maximize the probability of trial-by-trial shock expectancy or closeness updates given the specific model. Both the Learning Models and Closeness Models were estimated based on a linear likelihood function.

Parameters were independently fitted to each participant using the Broyden-Fletcher-Goldfarb-Shanno (BFGS) optimization method. To avoid local minima in parameter fitting, optimization was initiated with 5 randomly selected start values. Model implementations and parameter fitting was done in R-3.5.1 (R Development Core Team, 2012). To examine the degree to which the model explained the experimental data, we also calculated the mean squared error over the shock expectancy and closeness ratings.

We compared models with the Bayesian Information Criterion (BIC), which penalizes the model evidence with model complexity as follows: BIC = −2ln(L) + ln(n)k, where −ln(L) is the negative log-likelihood, n is the number of responses used to compute the likelihood, and k is the number of model parameters. BIC measures are summed across all participants. A smaller BIC indicates a better model fit.

### Linking parameter estimates to change in intergroup impressions

We performed multiple linear regression analyses in R-3.5.1 (R Development Core Team, 2012) to investigate whether the change in intergroup impressions was mainly driven by the unexpected experiences with outgroup members (as predicted by outgroup-focused theories), with the ingroup members (as predicted by ingroup-focused theories), or with both. The computational parameters (i.e., *W_ingroup_, W_outgroup_, γ_ingroup_, γ_outgroup_*) estimated from Closeness Model 1 (see details in the computational modeling section) were entered as predictors. In an additional analysis, the modern racism score was entered as a covariate to control for the influence of outgroup attitudes. The change of ingroup and outgroup impressions over the learning experiment was entered as the dependent variable (i.e. (ingroup-outgroup)_before_ – (ingroup-outgroup)_after_). Both the predictors and the dependent variable were standardized.

To examine whether ingroup identification moderated the associations between the weight given to ingroup prediction errors (*W_ingroup_*) and the intergroup impression change, we performed a moderation analysis with the *W_ingroup_* as the independent variable and individuals’ ingroup identification scores as the moderator. The independent variable and the moderator were standardized and multiplied together to generate the interaction term. Then the independent variable, moderator and the interaction term were entered into the regression model to associate with the intergroup impression change (i.e. (ingroup-outgroup)_before_ – (ingroup-outgroup)_after_). Again, we also performed an additional analysis in which the modern racism score was entered as a covariate to control for the influence of outgroup attitudes.

### Image acquisition and analyses

The experiment was conducted on a 3-T Philips whole-body MR scanner (Philips Medical Systems), equipped with an eight-channel Philips SENSitivity Encoded (SENSE) head coil. Structural image acquisition consisted of T1-weighted images (voxel size 1×1×1mm). For functional imaging, we used T2*-weighted echo-planar imaging (number of slices: 40, repetition time: 2.38s, voxel size: 3 x 3 x 3mm, field of view: 240 x 240mm, echo time: 30 ms, flip angle: 90).

Imaging data were analyzed in SPM12 (https://www.fil.ion.ucl.ac.uk/spm/software/spm12/). We followed a standardized preprocessing procedure. Functional images were first slice-time corrected, realigned and coregistered to the anatomical image of the participant. The anatomical image was processed using a unified segmentation procedure combining segmentation, bias correction, and spatial normalization to the MNI template (Ashburner & Friston, 2005); the same normalization parameters were then used to normalize the EPI images. Lastly, the functional images were spatially smoothed using an isotropic 8 mm full-width at half-maximum (FWHM) Gaussian kernel.

Whole-brain analyses were conducted to examine the neural correlates of prediction errors. The first level general linear model (GLM) analyses included the onsets and durations of fixation, closeness ratings, room revelation, outcome expectancy rating, feedback revelation, and shock delivery/omission. Parametric modulators modeled the prediction errors (derived from the reinforcement learning model for the model-derived analysis and derived from the difference between the outcome and participants’ shock expectancy ratings for the model-independent analysis) at the feedback revelation. The closeness ratings, room revelation and feedback revelation were modeled separately for the onsets of ingroup and outgroup trials. These regressors were convolved with the canonical hemodynamic response function and its time derivatives. Finally, the model contained six (three translation and three rotation) regressors to account for motion.

We assessed prediction error-related activity in a random effect model with one-sample t-tests on the contrast images created by the parametric modulators. We first analyzed prediction error-related activation independently of group membership. To do so, we weighted both ingroup and outgroup prediction errors regressors with a 1 on the first level and used the resulting contrast images to perform a one-sample t-test against zero on the second level. The regions encoding positive or negative prediction error were defined as the regions positively or negatively associated with prediction errors (i.e., significantly larger or smaller than zero) in the second level analysis (i.e., ingroup prediction errors + outgroup prediction errors). We also contrasted ingroup vs. outgroup prediction error-related activity at the first level and then compared the resulting images against zero in the second level analysis. Imaging results were determined in whole-brain analyses, using a combined voxel-level threshold of P_uncorrected_ < 0.001 and a family-wise error (FWE) corrected cluster-level threshold of P < 0.05.

### Psychophysiological interaction (PPI) analysis

To examine functional connectivity, we performed a psychophysiological interaction (PPI) analysis (Friston et al., 1997; McLaren et al., 2012), using the generalized PPI (gPPI) toolbox (https://www.nitrc.org/projects/gppi), in which the inclusion of task regressors reduces the possibility of detecting functional connectivity simply caused by co-activation (McLaren et al., 2012).We extracted the time series of the left IPL cluster (i.e. 5mm sphere centered on the peak voxel [x/y/z = −24/-55/53]), which showed differences between ingroup and outgroup prediction errors, as the physiological regressor. The condition onset times modelled in the analysis of activity (i.e., fixation, closeness rating, room revelation, outcome expectancy rating, feedback revelation, and shock delivery/omission) were separately convolved with a canonical hemodynamic response function, creating the psychological regressors. Then the physiological regressor was multiplied with the psychological regressors to obtain the psychophysiological interaction terms (PPI regressor). The physiological, psychological and psychophysiological interaction regressors as well as six motion parameters were entered into the GLM. We first used this GLM model to determine regions in which connectivity strength with the left IPL was modulated by revealing the ingroup or outgroup outcome (vs. the implicit baseline) at the first level analyses. Thus, we put a weight of 1 on the PPI regressor in which the corresponding psychological regressor was the onset time of in/out-group outcome revelation, and a weight of 0 on all other regressors at the first level. Next, we determined regions whose connectivity strength with the left IPL was modulated by the weight given to ingroup and outgroup prediction errors (W*_ingroup_*/W*_outgroup_*). To do so, we conducted second level analyses with the *W_ingroup_* and *W_outgroup_* as the predictors that were regressed against the contrast images of the PPI regressors acquired from the first level analyses.

To test the a-priori assumption that the individual *W_ingroup_* parameter alters the functional connectivity between the IPL (i.e., the region that showed an ingroup bias for the processing of negative prediction errors), and regions that process negative prediction errors, we performed small-volume correction in the regions encoding negative prediction errors (i.e. all regions where activity survived a combined voxel-level threshold of P_uncorrected_ < 0.001 and a family-wise error (FWE) corrected cluster-level threshold of P < 0.05 in previous whole-brain level analysis). Significant effects in the small-volume correction were reported using a combined voxel-level threshold of P_uncorrected_ < 0.001 and corrected cluster-level threshold of P < 0.05 based on a family-wise error (FWE) cluster-level correction.

To evaluate the correlation between the left IPL-AI coupling and the impression change, we extracted the connectivity strength for the bilateral AI regions identified by the PPI analysis (left AI: peak coordinate: −45/2/5; right AI: peak coordinate: 42/17/2; 5 mm sphere centered at the peak coordinate). The beta values of the ROI were extracted using MarsBaR (http://marsbar.sourceforge.net). The modern racism score was entered as covariate. We performed moderation analysis to test whether the association between the weight parameter (i.e. *W_ingroup_* and *W_outgroup_*) and the left IPL-bilateral AI coupling was significantly stronger when participants received feedback from the ingroup compared to the outgroup. Group (ingroup vs. outgroup) was entered as the moderator and coded as a dichotomous dummy variable (0 representing ingroup and 1 representing outgroup). Weight parameters and group were then standardized and multiplied to generate the interaction term. The independent variable, moderator and the interaction term were entered into the regression model to test the association with left IPL – bilateral AI coupling (i.e., connectivity strength extracted above).

## Results

### Behavioral evidence for impression change

To assess how learning affected in- and outgroup impressions, participants completed a standardized impression scale for the in- and outgroup before and after the learning experiment (see Methods for details). We conducted a linear mixed model (LMM) with group (ingroup/outgroup), time (before/after-learning) and group × time as predictors, and the rating scores on the impression scale as dependent variable. The results revealed a significant group × time interaction (*χ^2^*(1) = 4.44, *p* = 0.035, β = −0.37, SE = 0.18, **Figure 2a**). Clarifying the interaction effect, separate analyses showed significantly more positive impressions towards ingroup compared to outgroup members before learning (*χ^2^*(1) = 8.02, *p* = 0.005, β = −0.52, SE = 0.18), but no significant effect of group after learning (*χ^2^*(1) = 1.24, *p* = 0.26, β = −0.15, SE = 0.13).We also determined participants’ closeness ratings towards the ingroup and outgroup at the beginning (the first trial) and the end (the last trial) of the experiment. We observed a significant group × time interaction in closeness ratings (*χ^2^*(1) = 6.87, *p* = 0.009, *β* =1.44, SE = 0.55, **Figure 2b**). Elucidating the interaction effect, separate analyses revealed higher closeness ratings for the ingroup compared to the outgroup before learning (*χ^2^*(1) = 3.91, *p* = 0.048, β = −0.82, SE = 0.41), and a trend level of higher closeness for the outgroup compared to the ingroup after learning (*χ^2^*(1) = 2.86, *p* = 0.091, β = 0.62, SE = 0.37).

**Figure 2.**
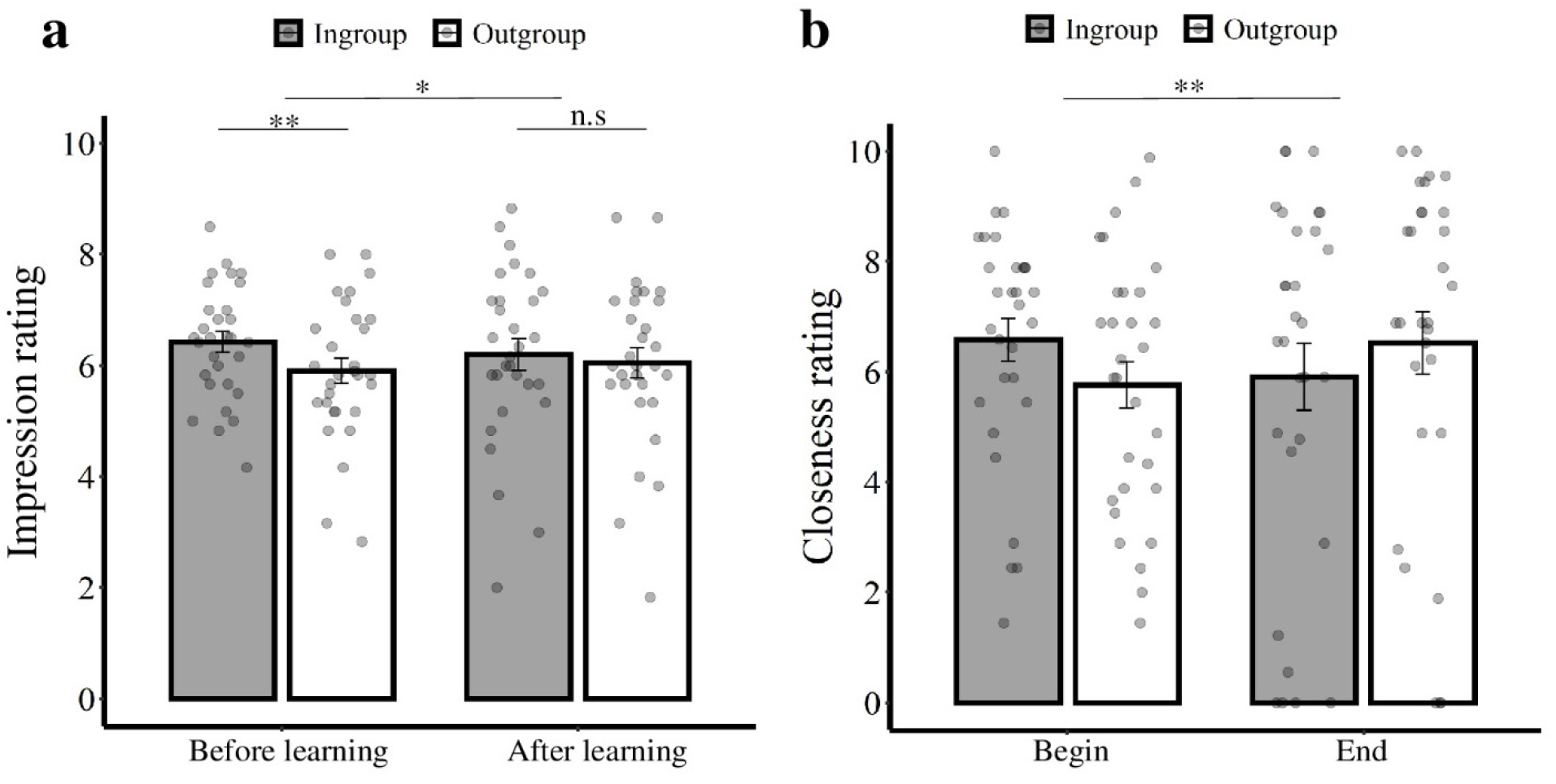
Impression and closeness ratings before and after learning. (a) Impression ratings before and after the learning experiment. Larger values represent more positive impressions. (b) Closeness ratings at the beginning and end of the experiment. For both types of ratings, the initial ingroup bias was reduced after learning. Error bars represent standard errors of the mean.

### Computational processes underlying dynamic changes in shock expectancy and ingroup and outgroup closeness ratings

To understand how the learning experience changed in- and outgroup impressions, we examined the computational processes underlying the dynamic changes in ratings. First, we modelled participants’ trial-by-trial shock expectancies using a Rescorla-Wagner reinforcement-learning model (Rescorla & Wagner, 1972) to test for differences in basic learning mechanisms between the ingroup and outgroup conditions. Model comparison revealed that shock expectancies concerning the ingroup and outgroup were best characterized by a model (Learning Model 3) with different learning rates and response parameters for ingroup and outgroup experiences (*r*^2^ = 0.29 ± 0.26; mean ± SD; **Figure 3a**, see **Table S1** and methods for details of model comparisons). The estimated learning rate was larger for the outgroup than for the ingroup, whereas the estimated response sensitivity parameters were comparable for the two groups (**Table S3**, *α: t*(29) = 2.30, *p* = 0.029, 95% CI = [0.005, 0.088]. β: *t*(29) = 0.770, *p* = 0.448, 95% CI = [-0.0 0.19]).

**Figure 3.**
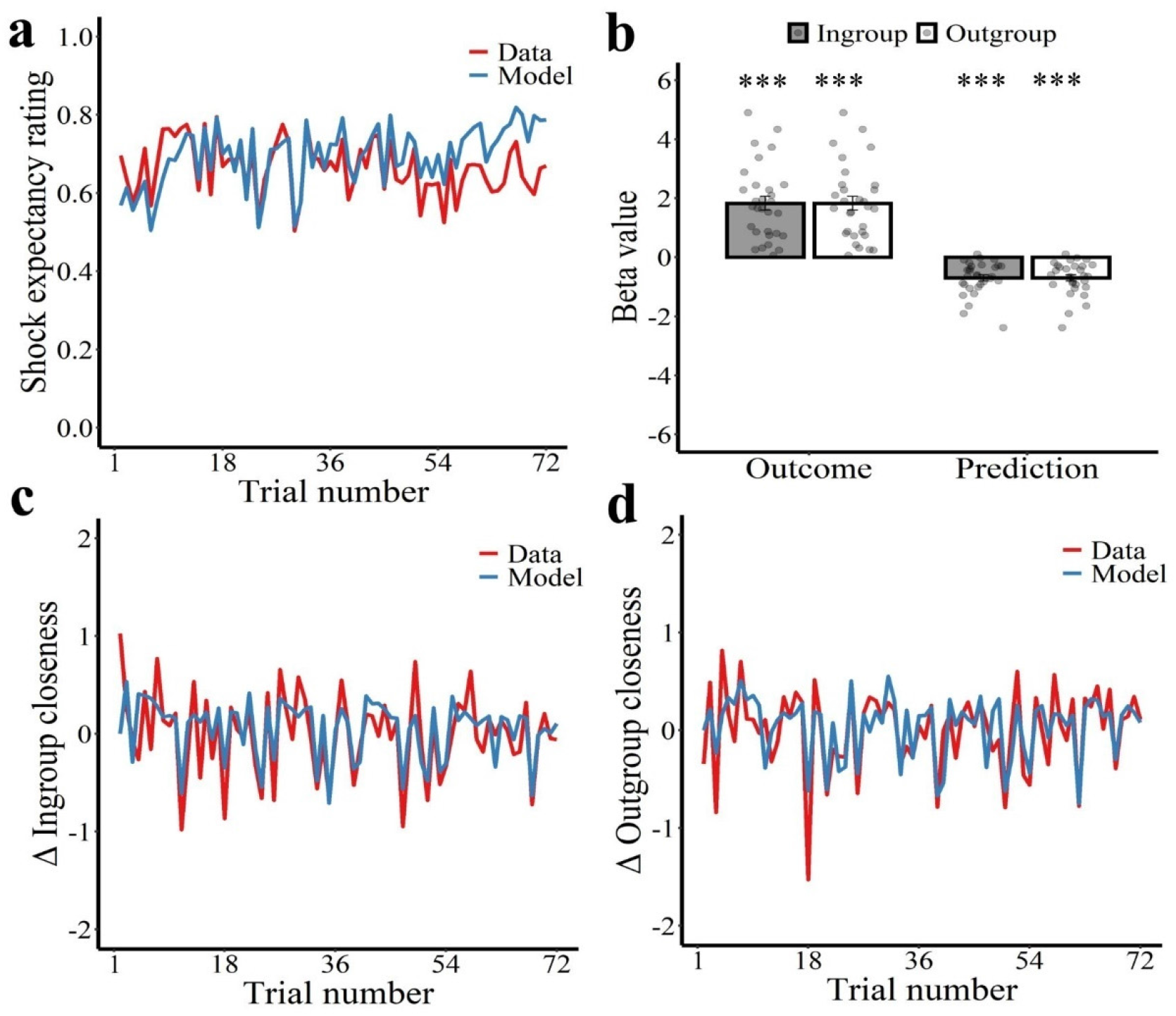
Computational models explain shock expectancy ratings and changes of in- and outgroup closeness ratings. (a) Predictions of receiving shocks (red line) varied over the course of the experiment and our learning model explained these predictions (blue line). The model estimates illustrate the best fitting model for shock expectancy ratings. (b) Results of linear mixed model analyses. Both experienced and expected outcome influenced change in ingroup and outgroup closeness ratings. \*\**P* < 0.001. Error bars represent standard errors of the mean. (c) Trial-by-trial changes of closeness ratings (red) and corresponding model estimates (blue) for the ingroup and (d) for the outgroup. The model estimates illustrate the best fitting Closeness model.

Next, we tested how ingroup and outgroup prediction errors (i.e., the difference between received feedback and expected feedback) affected trial-by-trial changes in ingroup and outgroup closeness ratings. As the ratings tended to be auto-correlated across trials, we used the trial-wise difference in closeness ratings to reduce the correlation (a standard method in time series analysis, Seth, 2010). Notably, the changes in closeness to the ingroup were only weakly correlated to the changes in closeness to the outgroup (average *r* across subjects = 0.14).

We first tested, in a model-independent manner, the hypothesis that dynamic changes in closeness ratings depend both on the nature of the outcome and the expectations about the outcome. If prediction errors explain changes in closeness better than the outcome alone, the trial wise updates of closeness should not only be positively correlated with the nature of the outcome (a positive outcome corresponds to positive experiences related to not receiving shock) but also negatively correlated with the expected outcome (Rutledge et al., 2014; Will et al., 2017). To this end, we first fitted an LMM to test the influence of experienced outcome and expected outcome on the changes of closeness. As hypothesized, for the changes of closeness towards both ingroup and outgroup, we found a significant positive correlation with experienced outcome (**Figure 3b**, ingroup: *χ^2^*(1) = 49.57, *p* < 1×10^-12^, *β* =1.83, SE = 0.26; outgroup: *χ^2^*(1) = 37.98, *p* < 1×10^-10^, β =1.65, SE = 0.27), but a negative association with expected outcome (**Figure 3b**, ingroup: *χ^2^*(1) = 10.55, *p* = 0.001, β = −0.70, SE = 0.22; outgroup: *χ^2^*(1) = 18.89, *p* < 1×10^-5^, β = −0.89, SE = 0.20). Thus, the two main constituents of prediction errors i.e., experienced and expected outcomes explain trial-by-trial changes in closeness ratings.

Next, we formally modeled the trial-by-trial update of in- and outgroup closeness ratings as a linear function of the cumulative impact of prediction errors, as estimated by the best fitting reinforcement learning model. Our winning closeness model (Equation 4, Closeness Model 1) successfully captured dynamic changes of closeness ratings at the individual level for both ingroup (*r^2^* = 0.19 ± 0.16; mean ± SD; see **Figure 3c**) and outgroup (*r^2^* = 0.21 ± 0.18; mean ± SD; see **Figure 3d**). The winning model assumes that the changes in closeness to group *i* (i.e., the ingroup or the outgroup) for each trial *t* are driven by the time-discounted sum of previous prediction errors to outcomes from the corresponding group *i*. We chose this model because it outperformed a range of alternatives including a model that solely considered experienced outcome, but not outcome expectations (see **Table S2** for details of model comparisons).

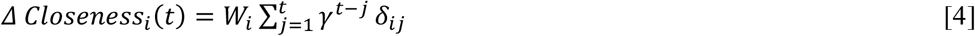

The winning model (Equation 4, Closeness Model 1) has two free parameters to capture the changes in closeness to each group, the weight *W* (−10 ≤ *W* ≤ 10, bounded by the rating scales), which determines the average influence of prediction errors (*δ_ij_*) arising from group *i* on changes in closeness towards group *i*, and the discount parameter *γ* (0 ≤ *γ*≤ 1), which determines how steeply older prediction errors are discounted (see Methods for additional details). If *γ* is close to one, all preceding prediction errors receive the same weight, and if it is close to zero, only the last prediction error leads to subsequent changes in closeness.

We fitted the changes in ingroup and outgroup closeness rating separately and estimated group-specific parameters (*W_ingroup_, W_outgroup_, γ_ingroup_, γ_outgroup_*). The parameters estimated by the winning model (the weight parameter *W* and the discount parameter *γ*), did not differ between the in- and outgroup (**Table S3**, *W*: *t*(29) = 0.28, *p* = 0.790, 95% CI = [−0.28, 0.36]; *γ*: t(29) = −0.16, *p* = 0.870, 95% CI = [-0.14, 0.12]). We also confirmed that prediction errors, rather than the experience of shock per se, gave the best account of the data (**Table S2**, Ingroup: ΔBIC = 88, Outgroup: ΔBIC = 179). Together, these results demonstrate that ingroup and outgroup closeness is dynamically updated based on prediction errors that are elicited by the behavior of in- and outgroup members.

### Comparing the influence of ingroup and outgroup prediction errors

So far, our results indicate that ingroup and outgroup experiences both generated prediction errors that resulted in learning-related updates of ingroup and outgroup closeness. Next, we investigated how the learning from ingroup and outgroup experiences relates to the changes in intergroup impressions shown in **Figure 2a**. To this end, we tested whether the observed update in impressions was mainly driven by the experiences with the outgroup members (as predicted by outgroup-focused theories), by experiences with the ingroup members (as predicted by ingroup-focused theories), or by both.

To address this question, we conducted a multiple linear regression analysis with the changes of ingroup and outgroup impressions before and after learning as dependent variable, i.e. (ingroup-outgroup)_before_ – (ingroup-outgroup)_after_. As independent variables we used the *W_ingroup_* and *W_outgroup_* parameters (reflecting the weight with which ingroup and outgroup prediction errors affect the updates in ingroup and outgroup closeness), as well as *γ_ingroup_* and *γ_outgroup_* parameters (reflecting how steeply older prediction errors are discounted). The variance inflation factor (VIF) for all variables in the regression was < 3.4, indicating that multicollinearity was not a concern. The results revealed that the reduction of the ingroup bias in impression ratings was uniquely predicted by the *W_ingroup_* parameter (β = 0.76, SE = 0.29, *t* = 2.72, *p* = 0.014, 95% CI [0.171 1.357], **Table 1, Figure 4a**). Importantly, the more strongly the individual weighted ingroup prediction errors, the more pronounced the reduction of ingroup bias in impression ratings was (**Figure 4b**). By contrast, the *W_outgroup_* parameter did not significantly predict the decline of ingroup favoritism (β = −0.16, SE = 0.30, *t* = −0.49, *p* = 0.608, 95% CI [-0.769 0.460], **Figure 4a**). These results show that a decrease of ingroup closeness due to a violation of ingroup predictions can result in a decline of ingroup favoritism.

**Figure 4.**
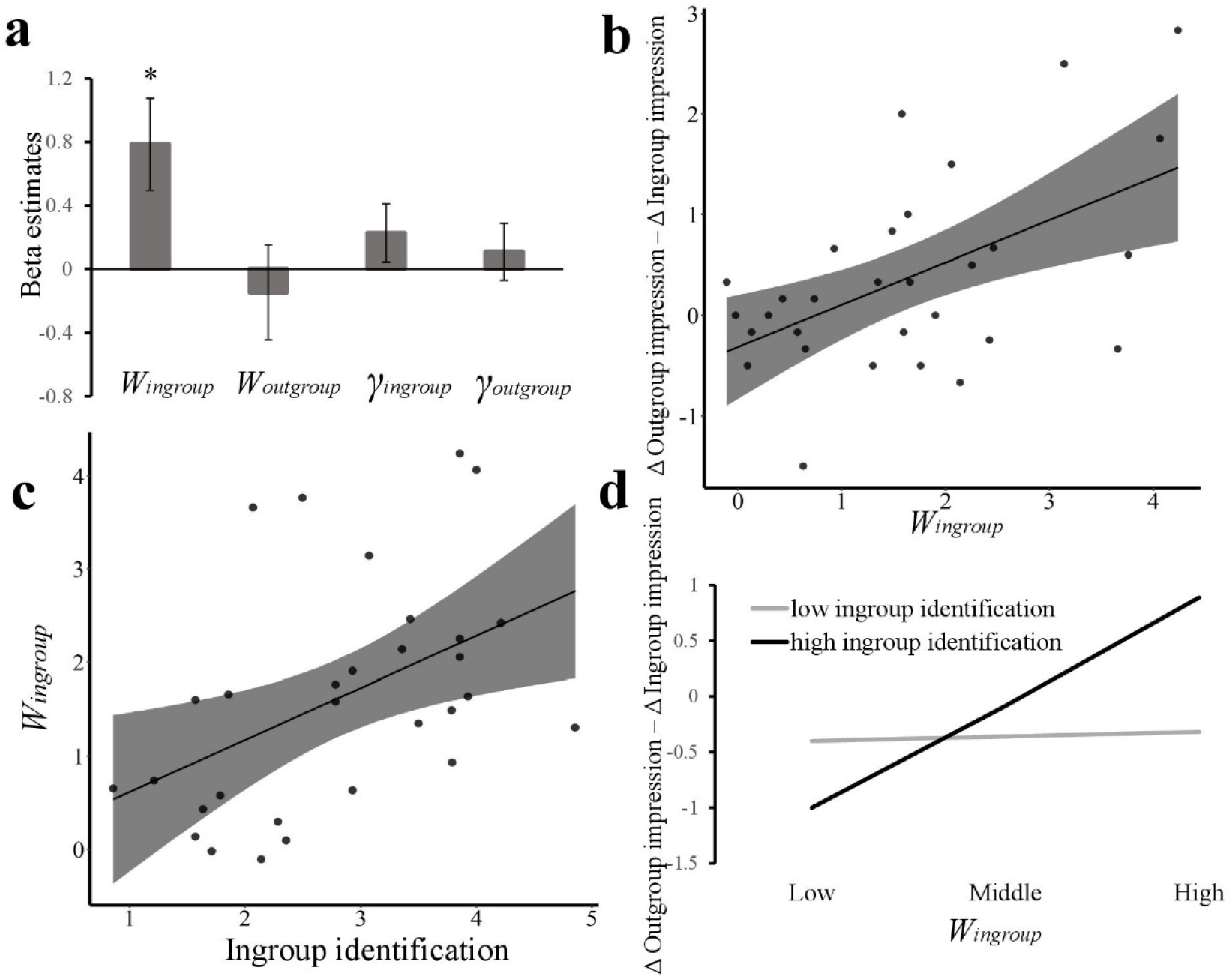
Behavioral results support ingroup focused theories of intergroup impression change. (a) Linear regression predicting impression change (Δoutgroup impression - Δingroup impression) with computational model parameters. The regression coefficient for *W_ingroup_* was significant, but that for *W_outgroup_* was not. Error bars represent standard errors of the mean. (b) Correlation between *W_ingroup_* parameter and decline of ingroup favoritism as measured by impression rating. (c) Correlation between ingroup identification and the *W_ingroup_* parameter. (d) Moderation analysis. Ingroup identification moderated the relationship between the *W_ingroup_* parameter and pre-vs. post learning changes in intergroup impressions. Compared to people with weak ingroup identification (M-1SD), people with strong ingroup identification (M + 1SD) showed a stronger association between *W_ingroup_* and impression change.

**Table 1.**
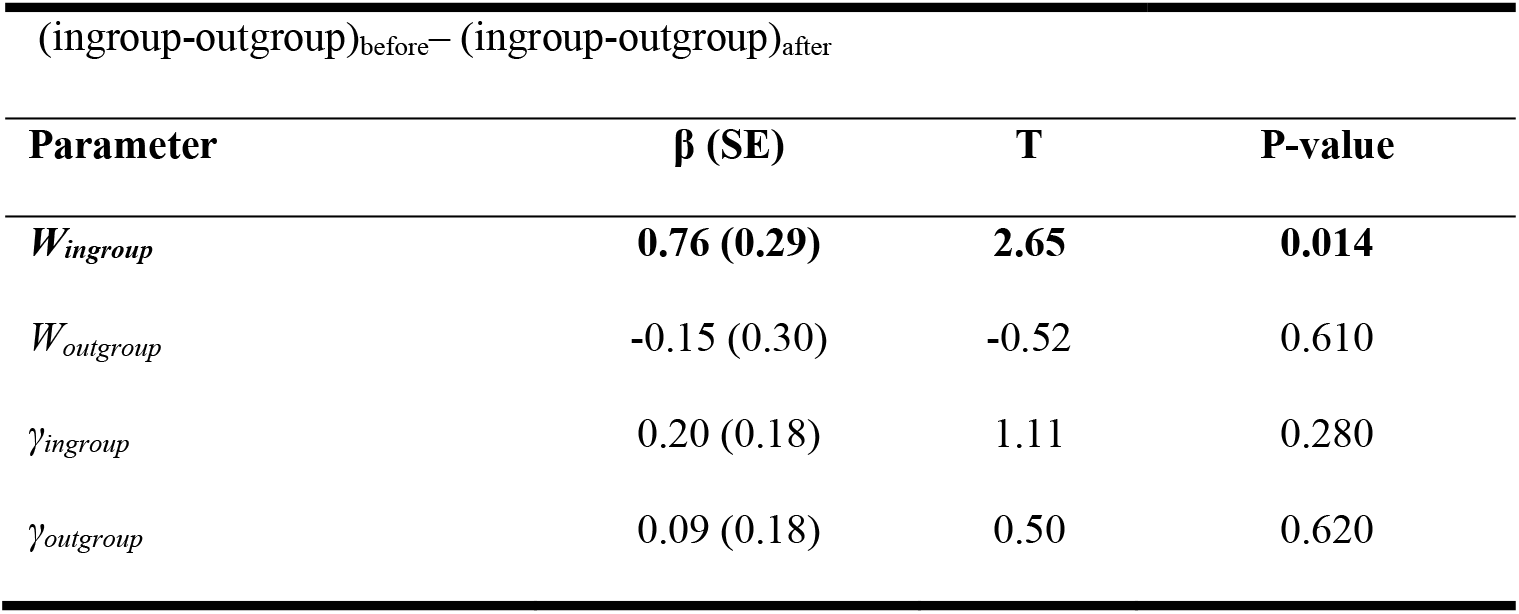
Predicting change in intergroup impressions from model parameters.

To confirm that *W_ingroup_*, predicted intergroup impression change irrespective of outgroup attitudes, we conducted the same regression analysis, using the individual scores in the modern racism scale as a covariate. Again, we found a significant effect for *W_ingroup_* (β = 1.01, SE = 0.28, *t* = 3.60, *p* = 0.001, 95% CI [0.433 1.595]), but not *W_outgroup_* (β = −0.41, SE = 0.29, *t* = −1.41, *p* = 0.172, 95% CI [-1.010 0.191]), suggesting that the positive relation between the weight given to ingroup prediction errors and reductions of ingroup favoritism were independent of outgroup attitudes.

To ascertain the specificity of these results, we also examined if the computational parameters estimated from the reinforcement learning model predict the changes of impressions. Neither the learning rates nor the response parameters for the ingroup or outgroup could predict the change of impressions (*p*s > 0.113). We also tested a regression model in which the *W_ingroup_, W_outgroup_, α_ingroup_* and *α_outgroup_* parameters were entered as independent variables to predict the impression change. Similarly, only the *W_ingroup_* was associated with the decline of ingroup favoritism (β = 0.69, SE = 0.31, *t* = 2.25, *p* = 0.034, 95% CI [0.058 1.316]), whereas the other variables were not related to the change of impressions (*p*s > 0.455). Therefore, the change of impression ratings was explained by the extent to which prediction errors influenced closeness ratings, and not by the prediction errors per se.

So far, the results of the regression analysis are in line with the ingroup-focused theories. According to one central prediction of the ingroup-focused theories, learning from ingroup experiences should be stronger in individuals with stronger ingroup identification, because they react more strongly to violations of ingroup expectations (Biernat et al., 1999; Branscombe et al., 1993; Mendoza et al., 2014). A correlation analysis between the individual scores of the ingroup identification scale (Leach et al., 2008) and the individual *W_ingroup_* and *W_outgroup_* parameter revealed a significant effect for *W_ingroup_* (*r* = 0.47, *p*_(Bonferroni-corrected)_ = 0.020, *p*_(uncorrected)_ = 0.010, 95% CI = [0.191, 0.701] **Figure 4c**), but not *W_outgroup_* (*r* = 0.28, *p*_(Bonferroni-corrected)_ = 0.274, *p*_(uncorrected)_ = 0.137, 95% CI = [-0.021 0.559]). The more a participant identified with the ingroup, the stronger the weight he gave to ingroup prediction errors. In contrast, individual differences in outgroup dislike (assessed with a modified version of the modern racism scale, McConahay,1986) were not related to either the *W_ingroup_* (*r* = −0.09, *p*_(uncorrected)_= 0.657, *p*_(Bonferroni-corrected)_ = 1, 95% CI [-0.379, 0.185]) or the *W_outgroup_* (*r* = 0.16, *p*_(uncorrected)_ 0.388, *p*_(Bonferroni-corrected)_ 0.776, 95% CI [-0.105, 0.422]) parameter.

To formally test the assumption that the relationship between *W_ingroup_* and intergroup impression change is moderated by ingroup identification, we conducted a moderation analysis with the ingroup identification scores as the moderator. *W_ingroup_* served as the independent variable and the intergroup impression change [(ingroup-outgroup)_before_ – (ingroup-outgroup)_after_] as the dependent variable. We found that the relationship between the *W_ingroup_* parameter and the pre-vs-post changes in intergroup impression ratings was significantly moderated by individual differences in ingroup identification (β = 0.34, SE = 0.19, *t* = 2.20, *p* = 0.037, **Figure 4d**). The more an individual identified with the ingroup, the stronger the relationship between *W_ingroup_* and intergroup impression change. The significant moderating effect of ingroup identification was also observed if we controlled for individual differences in outgroup attitudes by adding the responses on the modern racism scale as a covariate (β = 0.42, SE = 0.18, *t* = 2.30, *p* = 0.030). Taken together, both the regression analysis and the moderation analysis suggest that changes in intergroup impressions after intermittent ingroup and outgroup experiences are mainly associated with prediction error-related updates in ingroup closeness and with stronger effects in individuals that identify more strongly with the ingroup.

### Neural mechanisms underlying the ingroup-focused model

Having established that our computational model of dynamic changes in group identification can explain trial-by-trial changes in closeness and predict change in intergroup impressions, we then investigated the neural implementation of the model, by identifying brain activity corresponding to the estimated mechanisms and parameters. First, we investigated neural regions that encoded positive or negative prediction errors in general, i.e., irrespective of the group membership manipulation. To do so, we regressed the prediction error estimates against neural activity when the decision of the ingroup or outgroup individual was revealed. Negative prediction errors (i.e., shock prevention expected but shock delivered) were related to activity in dorsal anterior cingulate cortex (dACC) and anterior insula (AI). The more negative the prediction errors, the larger the neural responses in these regions. (**Figure 5a, Table 2**). Positive prediction errors (i.e., shock expected but prevented) correlated with activity in striatum, dorsomedial prefrontal cortex (dmPFC), ventromedial prefrontal cortex (vmPFC) and ventrolateral prefrontal cortex (vlPFC) (**Figure 5b, Table 2**). These classical learning regions showed stronger neural responses the more positive the prediction error.

**Figure 5.**
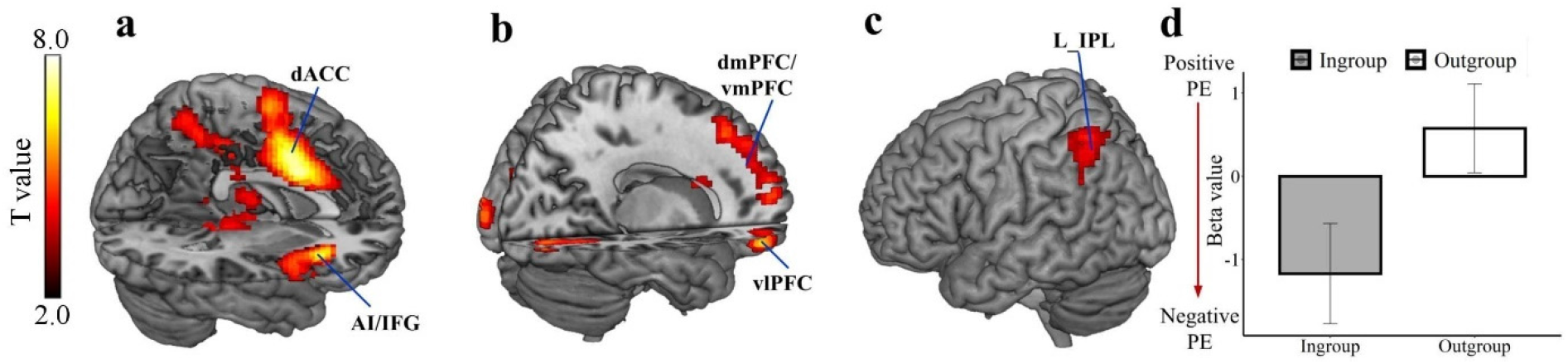
Neural activation related to the processing of prediction errors and ingroup versus outgroup group differences. (a) Negative prediction errors correlated with activity in dACC and AI. (b) Positive prediction errors correlated with activity in dmPFC, vmPFC, lPFC and striatum (caudate). (c and d) Left IPL activity was more strongly associated with negative prediction errors for the ingroup compared to the outgroup. Significant clusters were identified by combining a voxel-level threshold of *p* < .001 (uncorrected) and a cluster-level threshold of *p* < .05, FWE corrected across the whole brain. Display threshold at *p*_uncorrected_ < 0.001. Error bars represent standard errors of the mean.

**Table 2:**
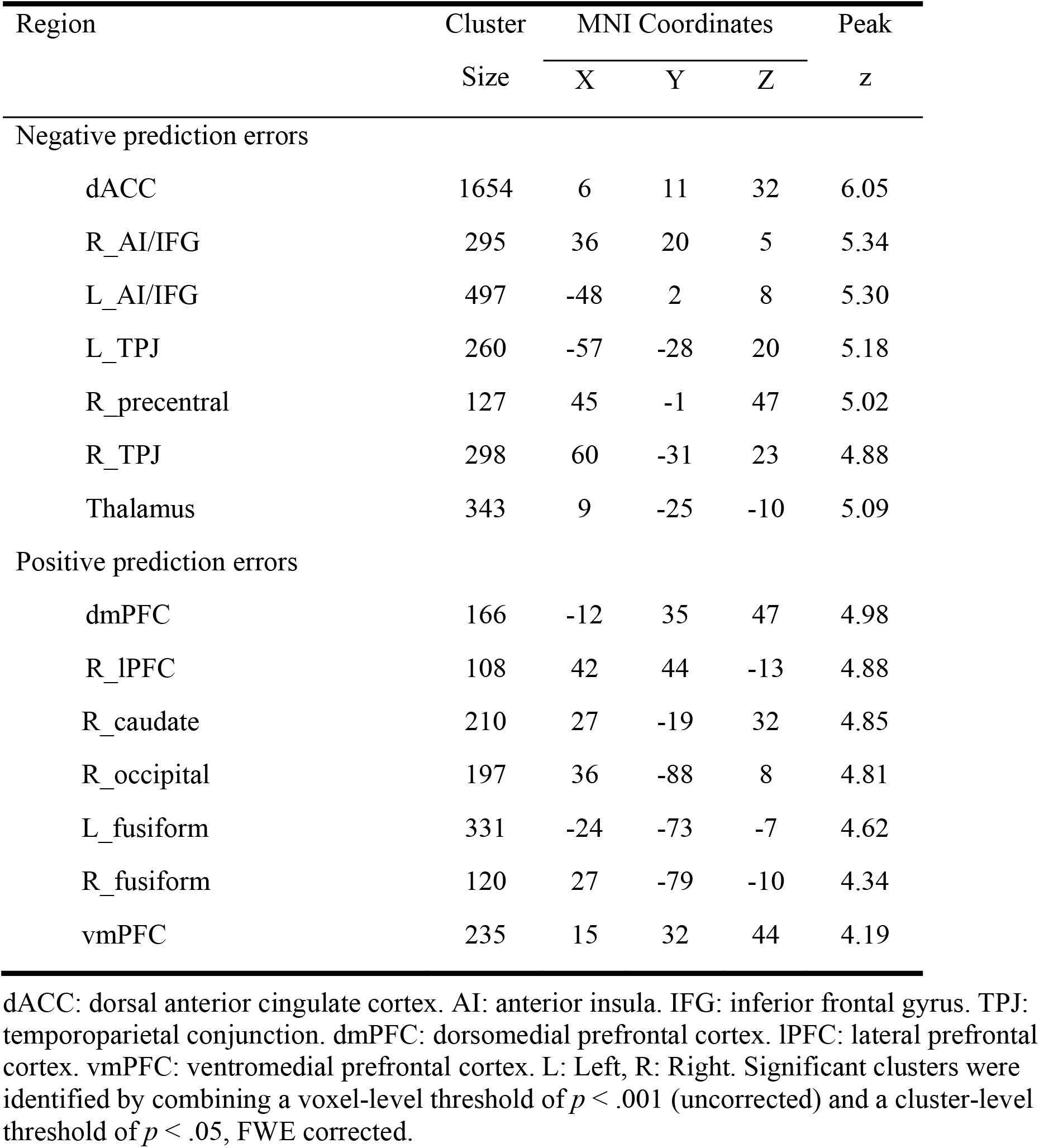
Brain regions that correlated with trial wise prediction errors.

Next, we contrasted the encoding of prediction errors between group conditions. We found a significant activation in the left inferior parietal lobule (IPL), indicating a stronger association with negative prediction errors for the ingroup compared to the outgroup (MNIxyz:-24/-55/53, Z_stats_= 4.19, *P*_FWE whole-brain corrected_ = 0.004) (**Figure 5c and 5d**). The reverse contrast did not show any significant results even at the very liberal threshold of *P*_uncorrected_ < 0.05 (whole-brain). A model-independent analysis, defining prediction errors as the difference between the outcome and participants’ shock expectancy ratings, revealed similar regions as those identified with model-derived prediction error analyses (**Figure S1**).

Our behavioral results suggest that the individual differences in the weight of the ingroup prediction error signal (*W_ingroup_*) predict the change in intergroup impressions. We reasoned that the weight given to the ingroup prediction errors could modulate the neural response in the IPL region that encoded ingroup prediction errors more strongly than outgroup prediction errors. However, a second level regression of the neural response elicited by ingroup prediction errors on the individual *W_ingroup_* parameters revealed no significant findings even at the very liberal threshold of P_uncorrected_ < 0.05 (whole-brain). Thus, ingroup prediction errors appear to be encoded independently of the individual weight of the ingroup prediction error.

Alternatively, it is possible that the individual *W_ingroup_* parameter alters the functional connectivity between the IPL (i.e., the region that showed an ingroup bias for the processing of negative prediction errors), and regions that process negative ingroup and outgroup prediction errors. We defined a region of interest (ROI) based on the entire neural network that was involved in the processing of negative prediction errors (**Table 2** and **Figure 5a**) and conducted a generalized psychophysiological interaction (gPPI) analysis (Friston et al., 1997; McLaren et al., 2012) to test if the connectivity between the IPL and regions encoding negative prediction errors is shaped by the individual *W_ingroup_* parameter. We estimated the connectivity strength between the left IPL (physiological regressor) and regions processing negative prediction errors when participants observed the ingroup feedback (psychological regressor, i.e., learned whether the ingroup individual was willing to help or not) in the first level analysis. We then conducted a second level regression analysis to test connections that were specifically associated with individual differences in *W_ingroup_* parameters after regressing out the influence of *W_outgroup_* parameters. Only connectivity between the left IPL and the left AI (MNIxyz:-45/2/5, Z_stats_= 3.96, *p* (SVC-FWE) = 0.022), and between the left IPL and the right AI (MNIxyz: 42/17/2, Z_stats_= 3.63, *p* (SVC-FWE) = 0.049) was modulated by *W_ingroup_* (**Figure 6a**). Specifically, individuals who weighted ingroup prediction errors more strongly to update the subjective closeness towards the ingroup (i.e., larger *W_ingroup_* parameters) also showed increased left IPL-AI coupling when receiving feedback from the ingroup.

**Figure 6.**
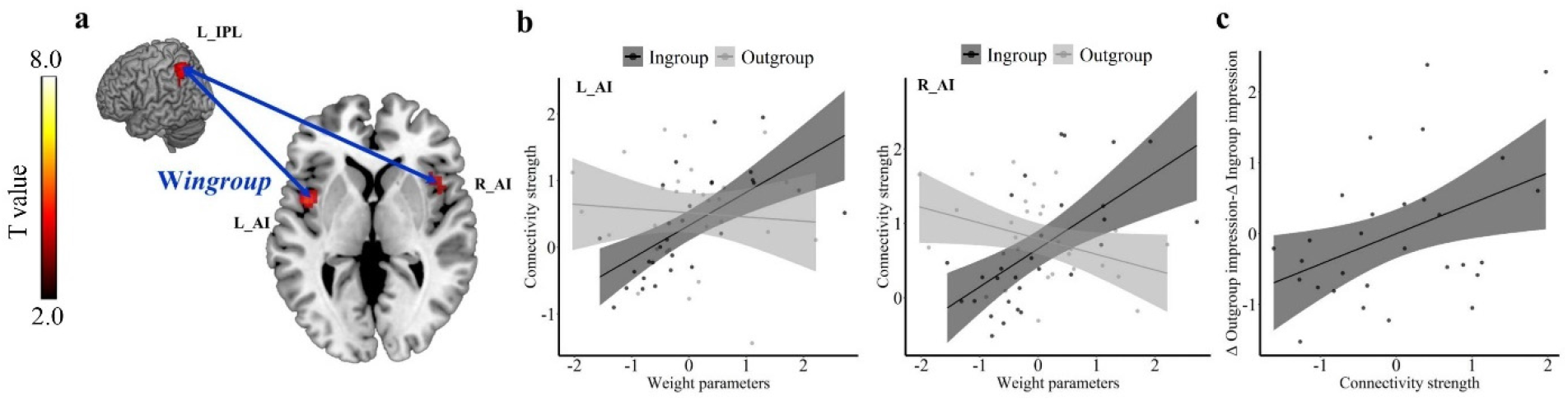
Results of the connectivity analysis. (a) The *W_ingroup_* parameter modulated the coupling between left IPL and bilateral AI during ingroup feedback revelation. Significant clusters were identified by SVC-FWE in the entire neural network that was involved in the processing of negative prediction errors shown in Figure 5a (b) The association between weight parameters and left IPL to bilateral AI coupling was significantly stronger when receiving feedback from ingroup compared to outgroup. (c) The connectivity strength between left IPL and left AI when receiving ingroup feedback correlated with the decline in ingroup bias in impression ratings. Display threshold at *p* < .001. Error bars represent standard errors of the mean.

To examine if this effect was specific for ingroup experience, we also conducted whole-brain level regression analyses with *W_outgroup_* as predictor (regressing out the influence of *W_ingroup_* parameters) on the connectivity strength between left IPL and the rest of the brain at the onset of outgroup feedback revelation, and found no significant regions even at lenient thresholds. To further validate this difference, we performed ROI-based moderation analyses and tested if group (i.e., ingroup vs. outgroup) significantly modulated the association between the weight parameters and IPL-AI connectivity strength. The moderation analysis showed that the interaction between weight parameters and group accounted for a significant proportion of variance in the connectivity strength between left IPL and bilateral AI at feedback onset (left AI: β = 0.36, SE = 0.12, *t* = 2.98, *p* = 0.004, right AI: β = 0.50, SE = 0.11, *t* = 4.42, *p* < 0.001, **Figure 6b**). Hence, the association between weight parameters and the strength of the connectivity between left IPL and bilateral AI was significantly stronger when participants received ingroup compared to outgroup feedback. These results corroborate our behavioral regression analysis, by demonstrating that the *W_ingroup_* parameters uniquely moderated the neural connectivity between left IPL and bilateral AI during ingroup feedback revelation and support the ingroup focused theories at the neural level.

Finally, as people who showed larger *W_ingroup_* also displayed greater decline of ingroup favoritism after learning, we tested whether the individual strength of the left IPL to bilateral AI connectivity during ingroup feedback revelation is related with the decrease of ingroup favoritism in impression rating. The modern racism score was entered as a covariate to control for individual differences in outgroup attitudes. Pearson correlation analyses showed that the connectivity strength between left IPL and left AI was positively associated with the reduction of the ingroup impression bias (*r* = 0.43, *p*_(Bonferroni-corrected)_ = 0.046, *p*_(uncorrected)_ = 0.023, 95% CI = [0.122, 0.645], **Figure 6c**). The correlation between the left IPL-right AI connectivity strength and intergroup impression change was not significant (*r* = 0.38, *p*_(Bonferroni-corrected)_ = 0.160, *p*_(uncorrected)_ = 0.080, 95% CI = [-0.089,0.651]). Moreover, there was no significant correlation between the left IPL-bilateral AI coupling and impression change when participants received feedback from the outgroup (left AI, *r* = 0.07, *p*_(Bonferroni-corrected)_ = 1, *p*_(uncorrected)_ = 0.729, 95% CI = [-0.223, 0.340]; right AI, *r* = 0.18, *p*_(Bonferroni-corrected)_ = 0.708, *p*_(uncorrected)_= 0.354, 95% CI = [-0.188, 0.513]). These results suggest that the neural connectivity between left IPL and left AI during the ingroup feedback was specifically related to the intergroup impression change.

## Discussion

We investigated how intermittent experiences with several ingroup and outgroup individuals dynamically change subjective closeness with, and impressions of, the ingroup and outgroup. Our results show that experiences with the ingroup can reduce ingroup favoritism (**Figure 2**), particularly when initial identification with the ingroup is strong. It is noteworthy that participants made identical experiences with the ingroup and outgroup, and fully processed the outgroup feedback (**Figure 3**), as indicated by comparable weights and discounting parameters for ingroup and outgroup prediction errors. The learning rate was even larger in the outgroup compared to the ingroup condition, suggesting that the expectancy of being shocked by the outgroup changed more easily based on outgroup experiences. However, outgroup learning rates did not predict changes in intergroup impressions. Instead, our results demonstrated that the intergroup impressions changed primarily as a result of ingroup experiences, i.e., the weight given to ingroup prediction errors (**Figure 4**).

The finding that ingroup experiences outweighed outgroup experiences, with regard to impression formation, supports ingroup-rather than outgroup-focused theories of group impression formation. In line with previous evidence showing an ingroup bias in information processing (Hughes et al., 2017; Howard & Rothbart, 1980; Foddy et al., 2009), our results show that learning from ingroup individuals can be more relevant for the formation of intergroup impressions compared to learning from outgroup individuals. Going beyond the existing evidence on ingroup biases in information processing (Hughes et al., 2017, Howard & Rothbart, 1980; Foddy et al., 2009), we reveal a mechanism through which ingroup experiences can affect intergroup impression formation. We demonstrate that intergroup impressions are shaped by learning-related changes in ingroup associations (closeness), caused by errors in the predictions of ingroup behavior. More generally, our findings show that models of impression formation can be enriched by learning processes originally established in the individual, non-social domain (Rescorla & Wagner, 1972), extending recent studies that investigated impression formation irrespective of intergroup processes (Mende-Siedlecki, 2018, Hackel et al., 2015, Ligneul et al., 2016).

On the neural level, the only region that showed a significant group difference (ingroup bias) in the processing of learning signals was the left IPL (**Figure 5c**). The IPL has been associated with the computation of “saliency maps” (Molenberghs et al., 2007; Vandenberghe & Gillebert, 2009; Kahnt et al., 2014), giving rise to biased internal representations of the external world (Treue, 2003). Moreover, there is evidence that the activity of the IPL is sensitive to group membership. For example, left IPL activity was stronger when participants perceived actions performed by ingroup compared to outgroup members, and this activation difference was further related to the individual ingroup bias in the judgment of action (Molenberghs et al., 2013). Extending previous findings, our results indicate that the IPL is also modulated by group membership in intergroup impression formation processes indexed by learning signals. Given its role in the computation of “saliency maps”, it is conceivable that the IPL increases the salience (i.e., the weight) of ingroup learning signals compared to outgroup learning signals. As such, intergroup impressions are more strongly formed by ingroup experiences, although participants learned from the ingroup and the outgroup feedback indicating that they paid attention to information from both social groups.

The individual extent of intergroup impression change was predicted by the individual extent of functional coupling between the IPL and the AI. The more strongly a person weighted the ingroup prediction errors, the stronger the coupling between the AI and the IPL, and the stronger the reduction in the ingroup bias with regard to impression ratings. The observed functional coupling between the left IPL and the AI is in line with previous findings of close structural connections between the AI and the parietal, frontal and occipital lobes, as well as limbic regions (Ghaziri et al., 2017). In particular, the IPL and AI are coactivated during motivationally consistent judgements (Leong et al., 2019) and learning of negative information generated by the ingroup (Hughes et al., 2017). Recent studies also highlighted the role of the AI in group categorization (Lieberman et al., 2005; Cikara et al., 2017), and in intergroup impression processing (Beer et al., 2008, Hein et al., 2016a; Hughes et al., 2017). For example, the AI has been shown to predict implicit negative attitudes towards members of a different race (Beer et al., 2008), and to contribute to learning-related impression change in intergroup contexts (Hein et al., 2016a; Hughes et al., 2017). Together, the current results suggest that the role of the AI in forming affective impressions is not limited to contexts with outgroup learning (Hein et al., 2016a), but extends to, and is facilitated by connectivity with IPL in contexts with ingroup learning.

In addition to its role in integrating social motivation and learning (Hein et al., 2016a), the AI is also a part of the “salience network”, which is particularly involved in the detection of motivationally salient stimuli, and is frequently associated with social emotion (Sridharan et al., 2008; Cauda et al., 2012; Craig & Craig, 2009; Amodio, 2014). Besides, the IPL has also been associated with salience processing and the computation of “saliency maps” (Molenberghs et al., 2007; Vandenberghe & Gillebert, 2009; Kahnt et al., 2014). Building on these findings, the present findings uncovered a connectivity between the IPL and the AI that might serve the selection of salient events (i.e. ingroup experiences) for enhanced processing of ingroup-specific information and subsequent adjustment of intergroup impressions.

In our study, participants made intermittent experiences with ingroup and outgroup individuals, and could therefore directly compare ingroup and outgroup experiences. In the context of direct group comparisons, group membership becomes more salient (Turner et al., 1987; Van Knippenberg & Wilke, 1988; Zhou et al., 2020). Given that individuals usually expect more positive outcomes from ingroup compared to outgroup individuals (Howard & Rothbart, 1980; Foddy et al., 2009), the equal experiences with the ingroup and the outgroup individuals did not meet these positive ingroup expectations, which may explain the higher relevance of ingroup prediction errors for intergroup impression change in our study.

Although our results emphasize the importance of ingroup experiences and ingroup associations for shaping intergroup impressions, they do not disregard the relevance of (positive) outgroup experiences. In line with previous studies (Hein et al., 2016a), our results highlight that experiences with outgroup individuals elicit learning-related signals (i.e., prediction errors) that change outgroup closeness. The outgroup-related learning processes had less impact on intergroup impressions than ingroup-related learning processes, but it is conceivable that they may have had a stronger effect in the absence of the more salient ingroup experiences.

## Conclusions

Our results demonstrated that intergroup impressions are mainly shaped by ingroup experiences, and revealed the underlying neural mechanisms. If their predictions about ingroup behavior are not confirmed by experience, people perceive outgroup individuals as positively as ingroup individuals. Based on this finding, portraying the ingroup in a more realistic light is a promising strategy to reduce ingroup favoritism, an insight that could have practical implications for the improvement of intergroup relations.

## Data availability

Behavioral data are available at: https://osf.io/ehqf4/.

Unthresholded statistical maps can be downloaded at https://neurovault.org/collections/11617/

## Code availability

Codes for modelling and analysis are available at: https://osf.io/ehqf4/.

## Authors’ Contributions

B.L., P.T. and G.H. conceived and designed the study. B.L. programmed the experiment. B.L., A.S. and P.K. collected the data. Y.Z. and B.L. analyzed the data with the input from G.H., P.T., A.S. and P.K. Y.Z, B.L. and G.H. drafted the manuscript and all authors edited the manuscript.

## Acknowledgements

This work was supported by the German Research Foundation (GH, HE 4566/5-1; HE 4566/3-1) to G.H, an Emmy Noether fellowship of the German Research Foundation (SO 1636/2-1) to A.S, and Swiss NSF grants 100014_165884, 100019_176016 and IZKSZ3_162109 to P.T.

## Competing interests

The authors declare no competing interests.

## Notes

### Competing Interest Statement

The authors have declared no competing interest.

## References

1 Tajfel, H., Billig, M. G., Bundy, R. P., & Flament, C. Social categorization and intergroup behaviour. Eur. J. Soc. Psychol. 1, 149–178 (1971).

2 Greenwald, A. G., Poehlman, T. A., Uhlmann, E. L., & Banaji, M. R. Understanding and using the Implicit Association Test: III. Meta-analysis of predictive validity. J. Pers. Soc. Psychol. 97, 17 (2009).

3 Amodio, D. M., & Cikara, M. The social neuroscience of prejudice. Annu. Rev. Psychol. 72, 439–469 (2021).

4 Hein, G., Engelmann, J. B., Vollberg, M. C., & Tobler, P. N. How learning shapes the empathic brain. Proc. Natl Acad. Sci. USA. 113, 80–85 (2016a).

5 Mende-Siedlecki, P. Changing our minds: the neural bases of dynamic impression updating. Curr. Opin. Psychol. 24, 72–76 (2018).

6 Siegel, J. Z., Mathys, C., Rutledge, R. B., & Crockett, M. J. Beliefs about bad people are volatile. Nat. Hum. Behav. 2, 750–756 (2018).

7 Hackel, L. M., Doll, B. B., & Amodio, D. M. Instrumental learning of traits versus rewards: dissociable neural correlates and effects on choice. Nat .Neurosci. 18, 1233–1235 (2015).

8 Tobler, P. N., O’Doherty, J. P., Dolan, R. J., & Schultz, W. Human neural learning depends on reward prediction errors in the blocking paradigm. J. Neurophysiol.. 95, 301–310 (2006).

9 Burke, C. J., Tobler, P. N., Baddeley, M., & Schultz, W. Neural mechanisms of observational learning. Proc. Natl Acad. Sci. USA. 107, 14431–14436 (2010).

10 Kang, P., Burke, C. J., Tobler, P. N., & Hein, G. Why we learn less from observing outgroups. J. Neurosci. 41, 144–152 (2021).

11 Lindström, B., Selbing, I., Molapour, T., & Olsson, A. Racial bias shapes social reinforcement learning. Psychol. Sci. 25, 711–719 (2014).

12 Will, G.-J., Rutledge, R. B., Moutoussis, M., & Dolan, R. J. Neural and computational processes underlying dynamic changes in self-esteem. Elife. 6, e28098 (2017).

13 Hackel, L. M., Berg, J. J., Lindström, B. R., & Amodio, D. M. Model-based and model-free social cognition: investigating the role of habit in social attitude formation and choice. Front. Psychol. 10, 25–92 (2019).

14 Hackel, L. M., & Zaki, J. Propagation of economic inequality through reciprocity and reputation. Psychol. Sci. 29, 604–613 (2018).

15 Pettigrew, T. F., & Tropp, L. R. A meta-analytic test of intergroup contact theory. J. Pers. Soc. Psychol. 90, 751 (2006).

16 Lindström, B., & Tobler, P. N. Incidental ostracism emerges from simple learning mechanisms. Nat. Hum. Behav. 2, 405–414 (2018).

17 Barlow, F. K., et al. The contact caveat: Negative contact predicts increased prejudice more than positive contact predicts reduced prejudice. Pers. Soc. Psychol. Bull. 38, 1629–1643 (2012).

18 Rescorla, R. A., & Wagner, A. R. In Classical Conditioning II: Current Research and Theory. (eds. Black, A. H. & Prokasy, W. F.). 64–99. (Appleton-Century-Crofts, 1972).

19 Hackel, L. M., & Amodio, D. M. Computational neuroscience approaches to social cognition. Curr. Opin. Psychol. 24, 92–97 (2018).

20 Dovidio, J. F., Gaertner, S. L., & Kawakami, K. Intergroup contact: The past, present, and the future. Group processes & intergroup relations. 6, 5–21 (2003).

21 Bettencourt, B. A., Brewer, M. B., Croak, M. R., & Miller, N. Cooperation and the reduction of intergroup bias: The role of reward structure and social orientation. J. Exp. Soc. Psychol. 28, 301–319 (1992).

22 Marcus-Newhall, A., Miller, N., Holtz, R., & Brewer, M. B. Cross-cutting category membership with role assignment: A means of reducing intergroup bias. Br. J. Soc. Psychol. 32, 125–146 (1993).

23 Schlueter, E., & Scheepers, P. The relationship between outgroup size and anti-outgroup attitudes: A theoretical synthesis and empirical test of group threat-and intergroup contact theory. Soc. Sci. Res. 39, 285–295 (2010).

24 Chu, D., & Griffey, D. The contact theory of racial integration: The case of sport. Sociology of Sport Journal. 2, 323–333 (1985).

25 Paluck, E. L. Reducing intergroup prejudice and conflict using the media: a field experiment in Rwanda. J. Pers. Soc. Psychol. 96, 574 (2009).

26 Malhotra, D., & Liyanage, S. Long-term effects of peace workshops in protracted conflicts. J. Confl. Resolut. 49, 908–924 (2005).

27 Dovidio, J. F., & Gaertner, S. L. In Affect, cognition and stereotyping. 167–193. (Elsevier, 1993).

28 Hein, G., Engelmann, J. B., & Tobler, P. N. Pain relief provided by an outgroup member enhances analgesia. Proc. Royal Soc. B. 285, 20180501 (2018).

29 Tajfel, H., & Turner, J. C. In The Social Psychology of Intergroup Relations. (eds. Austin, W. G. & Worchel, S.). 33–47. (Brooks/Cole, 1979).

30 Maass, A., Salvi, D., Arcuri, L., & Semin, G. R. Language use in intergroup contexts: The linguistic intergroup bias. J. Pers. Soc. Psychol. 57, 981 (1989).

31 Foddy, M., Platow, M. J., & Yamagishi, T. Group-based trust in strangers: The role of stereotypes and expectations. Psychol. Sci. 20, 419–422 (2009).

32 Hughes, B. L., Zaki, J., & Ambady, N. Motivation alters impression formation and related neural systems. Soc. Cogn. Affect. Neurosci. 12, 49–60 (2017).

33 Howard, J. W., & Rothbart, M. Social categorization and memory for in-group and out-group behavior. J. Pers. Soc. Psychol. 38, 301 (1980).

34 Hutchison, P., Abrams, D., Gutierrez, R., & Viki, G. T. Getting rid of the bad ones: The relationship between group identification, deviant derogation, and identity maintenance. J. Exp. Soc. Psychol. 44, 874–881 (2008).

35 Marques, J. M., Yzerbyt, V. Y., & Leyens, J. P. The “black sheep effect”: Extremity of judgments towards ingroup members as a function of group identification. Eur. J. Soc. Psychol. 18, 1–16 (1988).

36 Mendoza, S. A., Lane, S. P., & Amodio, D. M. For members only: ingroup punishment of fairness norm violations in the ultimatum game. Soc. Psychol. Pers. Sci. 5, 662–670 (2014).

37 Biernat, M., Vescio, T. K., & Billings, L. S. Black sheep and expectancy violation: Integrating two models of social judgment. Eur. J. Soc. Psychol. 29, 523–542 (1999).

38 Branscombe, N. R., Wann, D. L., Noel, J. G., & Coleman, J. In-group or out-group extemity: Importance of the threatened social identity. Pers. Soc. Psychol. Bull. 19, 381–388 (1993).

39 Ligneul, R., Obeso, I., Ruff, C. C., & Dreher, J.-C. Dynamical representation of dominance relationships in the human rostromedial prefrontal cortex. Curr. Biol. 26, 3107–3115 (2016).

40 Park, B., Fareri, D., Delgado, M., & Young, L. The role of right temporoparietal junction in processing social prediction error across relationship contexts. Soc. Cogn. Affect. Neurosci. 16, 772–781 (2021).

41 Berry, J. W., & Sam, D. L. In The Oxford handbook of multicultural identity. (eds. Benet-Martínez, V. & Hong, Y.-y.). 97. (Oxford University Press, 2014).

42 Leach, C. W., et al. Group-level self-definition and self-investment: a hierarchical (multicomponent) model of in-group identification. J. Pers. Soc. Psychol. 95, 144 (2008).

43 McConahay, J. B. In Prejudice, discrimination, and racism. (eds. Dovidio, J. F. & Gaertner, S. L.). 91–125. (Academic Press, 1986).

44 Mende-Siedlecki, P., Cai, Y., & Todorov, A. The neural dynamics of updating person impressions. Soc. Cogn. Affect. Neurosci. 8, 623–631 (2013).

45 Sheng, F., Liu, Q., Li, H., Fang, F., & Han, S. Task modulations of racial bias in neural responses to others’ suffering. NeuroImage. 88, 263–270 (2014).

46 Wang, C., Wu, B., Liu, Y., Wu, X., & Han, S. Challenging emotional prejudice by changing self-concept: priming independent self-construal reduces racial in-group bias in neural responses to other’s pain. Soc. Cogn. Affect. Neurosci. 10, 1195–1201 (2015).

47 Dijksterhuis, A., & Van Knippenberg, A. The relation between perception and behavior, or how to win a game of trivial pursuit. J. Pers. Soc. Psychol. 74, 865 (1998).

48 Hein, G., Morishima, Y., Leiberg, S., Sul, S., & Fehr, E. The brain’s functional network architecture reveals human motives. Science. 351, 1074–1078 (2016b).

49 Hein, G., Silani, G., Preuschoff, K., Batson, C. D., & Singer, T. Neural responses to ingroup and outgroup members’ suffering predict individual differences in costly helping. Neuron. 68, 149–160 (2010).

50 Bates, D., Kliegl, R., Vasishth, S., & Baayen, H. Parsimonious mixed models. arXiv preprint arXiv:1506.04967. (2015).

51 Seth, A. K. A MATLAB toolbox for Granger causal connectivity analysis. J. Neurosci. Methods. 186, 262–273 (2010).

52 Ashburner, J., & Friston, K. J. Unified segmentation. NeuroImage. 26, 839–851 (2005).

53 Friston, K., et al. Psychophysiological and modulatory interactions in neuroimaging. NeuroImage. 6, 218–229 (1997).

54 McLaren, D. G., Ries, M. L., Xu, G., & Johnson, S. C. A generalized form of context-dependent psychophysiological interactions (gPPI): a comparison to standard approaches. NeuroImage. 61, 1277–1286 (2012).

55 Rutledge, R. B., Skandali, N., Dayan, P., & Dolan, R. J. A computational and neural model of momentary subjective well-being. Proc. Natl Acad. Sci. USA. 111, 12252–12257 (2014).

56 Molenberghs, P., Mesulam, M. M., Peeters, R., & Vandenberghe, R. R. Remapping attentional priorities: differential contribution of superior parietal lobule and intraparietal sulcus. Cereb. Cortex. 17, 2703–2712 (2007).

57 Vandenberghe, R., & Gillebert, C. R. Parcellation of parietal cortex: convergence between lesion-symptom mapping and mapping of the intact functioning brain. Behav. Brain Res. 199, 171–182 (2009).

58 Kahnt, T., Park, S. Q., Haynes, J.-D., & Tobler, P. N. Disentangling neural representations of value and salience in the human brain. Proc. Natl Acad. Sci. USA. 111, 5000–5005 (2014).

59 Treue, S. Visual attention: the where, what, how and why of saliency. Curr. Opin. Neurol. 13, 428–432 (2003).

60 Molenberghs, P., Halász, V., Mattingley, J. B., Vanman, E. J., & Cunnington, R. Seeing is believing: Neural mechanisms of action–perception are biased by team membership. Hum. Brain Mapp. 34, 2055–2068 (2013).

61 Ghaziri, J., et al. The corticocortical structural connectivity of the human insula. Cereb. Cortex. 27, 1216–1228 (2017).

62 Leong, Y. C., Hughes, B. L., Wang, Y., & Zaki, J. Neurocomputational mechanisms underlying motivated seeing. Nat. Hum. Behav. 3, 962–973 (2019).

63 Lieberman, M. D., Hariri, A., Jarcho, J. M., Eisenberger, N. I., & Bookheimer, S. Y. An fMRI investigation of race-related amygdala activity in African-American and Caucasian-American individuals. Nat .Neurosci. 8, 720–722 (2005).

64 Cikara, M., Van Bavel, J. J., Ingbretsen, Z. A., & Lau, T. Decoding “us” and “them”: Neural representations of generalized group concepts. J. Exp. Psychol. Gen. 146, 621 (2017).

65 Beer, J. S., et al. The Quadruple Process model approach to examining the neural underpinnings of prejudice. NeuroImage. 43, 775–783 (2008).

66 Sridharan, D., Levitin, D. J., & Menon, V. A critical role for the right fronto-insular cortex in switching between central-executive and default-mode networks. Proc. Natl Acad. Sci. USA. 105, 12569–12574 (2008).

67 Cauda, F., et al. Meta-analytic clustering of the insular cortex: characterizing the meta-analytic connectivity of the insula when involved in active tasks. NeuroImage. 62, 343–355 (2012).

68 Craig, A. D., & Craig, A. How do you feel--now? The anterior insula and human awareness. Nat. Rev. Neurosci. 10 (2009).

69 Amodio, D. M. The neuroscience of prejudice and stereotyping. Nat. Rev. Neurosci. 15, 670–682 (2014).

70 Turner, J. C., Hogg, M. A., Oakes, P. J., Reicher, S. D., & Wetherell, M. S. Rediscovering the social group: A self-categorization theory (Basil Blackwell, 1987).

71 Van Knippenberg, A., & Wilke, H. Social categorization and attitude change. Eur. J. Soc. Psychol. 18, 395–406 (1988).

72 Zhou, Y., et al. Neural dynamics of racial categorization predicts racial bias in face recognition and altruism. Nat. Hum. Behav. 4, 69–87 (2020).

