## Supplementary material for "Learning from ingroup experiences changes intergroup impressions": Figure S1; Table S1-S9

\# contributed equally to this work

* shared senior authorship

**Supplementary information**

**Figure S1**

**Table S1-9**

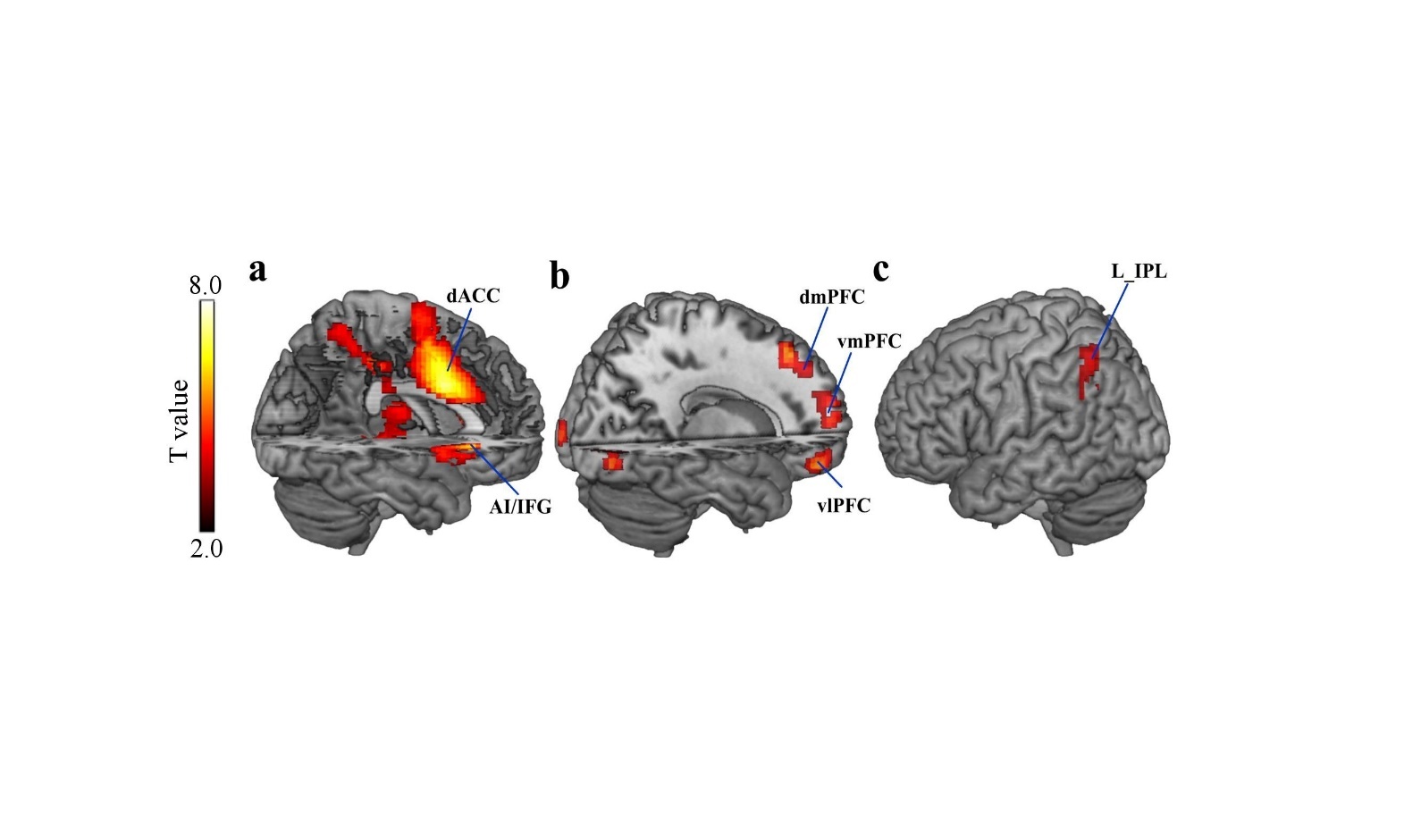

**Figure S1.** **Neural activation related to model-independent prediction errors.** Prediction errors were defined by the difference between the outcome and participants’ shock expectancy ratings. (a) Negative prediction errors correlated with activity in dACC and AI. (b) Positive prediction errors correlated with activity in striatum (caudate) dmPFC, vmPFC and lPFC. (c and d) left IPL activity was more strongly associated with negative prediction errors in the ingroup compared to the outgroup condition. Display threshold at *p* < .001, voxel > 60.

**Table S1.** **Comparisons of model fits for shock expectancy ratings.** Bayesian Information Criterion (BIC) measures are summed across all participants. Lower BIC values indicate better model fit. Mean squared error over shock expectancy indicate goodness of fit. K is the number of free parameters in each model. The winning model is shown in bold and italic.

| **Computational models** | **K** | **Mean r^2^** | **BIC** |
| --- | --- | --- | --- |
| **Shock expectancy ratings** |  |  |  |
| Learning Model 1: Same learning rate | 2 | 0.23 | -7638 |
| Learning Model 2: Two learning rates, one response parameter | 3 | 0.26 | -7664 |
| ***Learning Model 3: Two learning rates, two response parameters*** | ***4*** | ***0.29*** | ***-7717*** |
| Learning Model 4: Four learning rates, specific to both group and outcome | 5 | 0.25 | -6632 |

**Table S2.** **Comparisons of model fits for closeness ratings.** Bayesian Information Criterion (BIC) measures are summed across all participants. Lower BIC values indicate better model fit. Mean squared error over change of closeness ratings to ingroup or outgroup indicate goodness of fit. K is the number of free parameters in each model. The winning models are shown in bold and italic.

| **Computational models** | **K** | **Mean r^2^** | **BIC** |
| --- | --- | --- | --- |
| **Ingroup closeness ratings** |  |  |  |
| ***Closeness Model 1: Group-specific prediction error weights*** | ***2*** | ***0.19*** | ***220*** |
| Closeness Model 2: Outcome-specific prediction error weights | 3 | 0.21 | 282 |
| Closeness Model 3: Prediction error weights from both groups | 3 | 0.20 | 314 |
| Closeness Model 4: Experienced outcome only | 2 | 0.19 | 308 |
| **Outgroup closeness ratings** |  |  |  |
| ***Closeness Model 1: Group-specific prediction error weights*** | ***2*** | ***0.21*** | ***88*** |
| Closeness Model 2: Outcome-specific prediction error weights | 3 | 0.22 | 171 |
| Closeness Model 3: Prediction error weights from both groups | 3 | 0.21 | 216 |
| Closeness Model 4: Experienced outcome only | 2 | 0.17 | 267 |

**Table S3.** **Means and standard deviations for computational model parameters of the winning closeness model.**

| **Computational Parameters** | **Mean (SD)** |
| --- | --- |
| Weight on prediction error from ingroup (*W_ingroup_* ) | 1.63 (1.23) |
| Weight on prediction error from ingroup (*W_outgroup_*) | 1.59 (1.43) |
| Discounting factor for ingroup (*γ_ingroup_*) | 0.10 (0.26) |
| Discounting factor for outgroup (*γ_outgroup_*) | 0.11 (0.25) |
| Learning rate for ingroup (*α_ingroup_*) | 0.13 (0.12) |
| Learning rate for outgroup (*α_outgroup_*) | 0.17 (0.13) |
| Response parameter for ingroup (*β_ingroup_*) | 0.91 (0.39) |
| Response parameter for outgroup (*β_outgroup_*) | 0.96 (0.44) |

**Table S4.** **Impression and closeness rating change based on 29 participants.**

|  | Regressor | χ2 | p | β (SE) |
| --- | --- | --- | --- | --- |
| Impression rating | Group×Time | 3.93 | 0.047 | -0.36 (0.18) |
|  | Group (before learning) | 6.75 | 0.009 | -0.49 (0.19) |
|  | Group (after learning) | 0.85 | 0.360 | -0.13 (0.14) |
| Closeness rating | Group×Time | 5.77 | 0.016 | 1.30 (0.54) |
|  | Group( before learning) | 4.34 | 0.037 | -0.92 (0.44) |
|  | Group( after learning) | 2.60 | 0.110 | 0.65 (0.40) |

**Table S5.** **Comparisons of model fits for shock expectancy ratings based on 29 participants.** Bayesian Information Criterion (BIC) measures are summed across all participants. Lower BIC values indicate better model fit. Mean squared error over shock expectancy indicate goodness of fit. K is the number of free parameters in each model. The winning model is shown in bold and italic.

| **Computational models** | **K** | **Mean r^2^** | **BIC** |
| --- | --- | --- | --- |
| **Shock expectancy ratings** |  |  |  |
| Learning Model 1: Same learning rate | 2 | 0.23 | -7318 |
| Learning Model 2: Two learning rates, one response parameter | 3 | 0.26 | -7346 |
| ***Learning Model 3: Two learning rates, two response parameters*** | ***4*** | ***0.29*** | ***-7398*** |
| Learning Model 4: Four learning rates, specific to both group and outcome | 5 | 0.26 | -6412 |

**Table S6.** **Comparisons of model fits for closeness ratings based on 29 participants.** Bayesian Information Criterion (BIC) measures are summed across all participants. Lower BIC values indicate better model fit. Mean squared error over change of closeness ratings to ingroup or outgroup indicate goodness of fit. K is the number of free parameters in each model. The winning models are shown in bold and italic

| **Computational models** | **K** | **Mean r^2^** | **BIC** |
| --- | --- | --- | --- |
| **Ingroup closeness ratings** |  |  |  |
| ***Closeness Model 1: Group-specific prediction error weights*** | ***2*** | ***0.19*** | ***349*** |
| Closeness Model 2: Outcome-specific prediction error weights | 3 | 0.21 | 407 |
| Closeness Model 3: Prediction error weights from both groups | 3 | 0.20 | 439 |
| Closeness Model 4: Experienced outcome only | 2 | 0.18 | 429 |
| **Outgroup closeness ratings** |  |  |  |
| ***Closeness Model 1: Group-specific prediction error weights*** | ***2*** | ***0.21*** | ***93*** |
| Closeness Model 2: Outcome-specific prediction error weights | 3 | 0.22 | 176 |
| Closeness Model 3: Prediction error weights from both groups | 3 | 0.21 | 215 |
| Closeness Model 4: Experienced outcome only | 2 | 0.17 | 271 |

**Table S7.** **Means and standard deviations for computational model parameters of the winning model based on 29 participants.**

| **Computational Parameter** | **Mean (SD)** |
| --- | --- |
| Weight on prediction error from ingroup (*W_ingroup_* ) | 1.65 (1.25) |
| Weight on prediction error from ingroup (*W_outgroup_*) | 1.60 (1.46) |
| Discounting factor for ingroup (*γ_ingroup_*) | 0.10 (0.26) |
| Discounting factor for outgroup (*γ_outgroup_*) | 0.11 (0.25) |
| Learning rate for ingroup (*α_ingroup_*) | 0.12 (0.11) |
| Learning rate for outgroup (*α_outgroup_*) | 0.17 (0.13) |
| Response parameter for ingroup (*β_ingroup_*) | 0.93 (0.40) |
| Response parameter for outgroup (*β_outgroup_*) | 0.98 (0.44) |

**Table S8.** **Predicting change in intergroup impressions from model parameters based on 29 participants.**

| (ingroup-outgroup)_before_– (ingroup-outgroup)_after_ | |  |  |
| --- | --- | --- | --- |
| **Parameter** | **β (SE)** | **T** | **P-value** |
| ***W_ingroup_*** | ***0.79 (0.29)*** | ***2.72*** | ***0.012*** |
| *W_outgroup_* | -0.15 (0.30) | -0.49 | 0.630 |
| *γ_ingroup_* | 0.23 (0.18) | 1.23 | 0.230 |
| *γ_outgroup_* | 0.11 (0.18) | 0.60 | 0.550 |

**Table S9.** **Correlation and moderation analyses based on 29 participants.**

| **Correlation analyses** | r | *p*_(uncorrected)_ | *p*_(Bonferroni-corrected)_ |
| --- | --- | --- | --- |
| *W_ingroup_* & Ingroup identification | 0.495 | 0.006 | 0.012 |
| *W_outgroup_* & Ingroup identification | 0.296 | 0.119 | 0.238 |
| *W_ingroup_* & modern racism | -0.079 | 0.682 | 1 |
| *W_outgroup_* & modern racism | 0.168 | 0.384 | 0.768 |
| **Moderation analysis** | β (SE) | T | P-value |
| *W_ingroup_**Ingroup identification | 0.47 (0.20) | 2.39 | 0.025 |
